# Matrix factorization and transfer learning uncover regulatory biology across multiple single-cell ATAC-seq data sets

**DOI:** 10.1101/2020.01.30.927129

**Authors:** Rossin Erbe, Michael D. Kessler, Alexander V. Favorov, Hariharan Easwaran, Daria A. Gaykalova, Elana J. Fertig

## Abstract

While single-cell ATAC-seq analysis methods allow for robust clustering of cell types, the question of how to integrate multiple scATAC-seq data sets and/or sequencing modalities is still open. We present an analysis framework that enables such integration by applying the CoGAPS Matrix Factorization algorithm and the projectR transfer learning program to identify common regulatory patterns across scATAC-seq data sets. Using publicly available scATAC-seq data, we find patterns that accurately characterize cell types both within and across data sets. Furthermore, we demonstrate that these patterns are both consistent with current biological understanding and reflective of novel regulatory biology.

## Background

The Assay for Transposase Accessible Chromatin (ATAC-seq) subjects DNA to a hyperactive transposase in order to tag euchromatic regions of the genome for sequencing. ATAC-seq thus provides a quantitative estimate of genome-wide chromatin accessibility, and can be used to infer which genomic regions are most likely to interact directly with proteins and other biologically relevant molecules (1), (2). Specifically, accessibility at enhancers and promoters has considerable influence on the binding of transcription factors (TFs) and other transcriptional machinery (3). Quantification of accessibility at these regions enables the characterization of the regulatory biology that defines cell types and samples of interest (1), (2).

ATAC-seq data is often summarized by binning reads into data-defined genomic regions of frequent accessibility (generally termed peaks) or by aggregating the reads that contain annotated DNA motifs (e.g. transcription factor binding sites), which are collectively the targets of defined trans-acting factors (e.g. transcription factors) (4). Aggregating reads in these ways allows for a comparison of accessibility variation between samples and inference of the chromatin landscape of cell populations. However, the functional annotations available for these features are often incomplete, which can present significant challenges in the interpretation of ATAC-seq data, and can limit the integration of accessibility information across data sets. Furthermore, the high dimensionality and extreme sparsity of single cell ATAC-seq data (scATAC-seq) significantly compounds these analytic challenges, and further limits interpretation (5).

Therefore, computational methods are necessary to determine the patterns of accessibility that differentiate the regulatory biology associated with disparate cell populations in scATAC-seq data. Current tools for scATAC-seq analysis robustly cluster and annotate cell types. For example, ChromVAR, BROCKMAN, *Cusanovitch2018*, and scABC (6), (7), (8), (9) all output both clustering and inferred transcription factor binding within clusters, using clustering accuracy as their primary metric to evaluate efficacy. SnapATAC and cisTopic additionally provide the ability to query upregulated pathways from scATAC-seq data, but are still most strongly oriented towards the goal of effectively differentiating cell populations (5), (10). These methods provide effective tools for the analysis of individual scATAC-seq data, but require further extension to integrate the information learned from multiple scATAC-seq experiments or multiple sequencing modalities.

We develop a framework to enable cross-study and cross-platform analysis of multiple scATAC-seq data sets through the application of the Bayesian Non-Negative Matrix Factorization algorithm, CoGAPS, (11), (12) in conjunction with the transfer learning program projectR (13), (14). We demonstrate that CoGAPS simultaneously identifies robust cell types, upregulated pathways, and TF activity from scATAC-seq data. Notably, the projectR transfer learning method allows for the identification of the learned signatures of regulatory biology that we identify with CoGAPS within other datasets. Finally, we use matched RNA-seq data to provide orthogonal evidence for candidate regulatory mechanisms identified by our scATAC-seq analysis method. This workflow facilitates the development of consensus accessibility signatures for cellular populations using multiple data sets and data modalities. Furthermore, we demonstrate that combined CoGAPS analysis of scATAC-seq and scRNA-seq identifies novel biology, such as the association of the transcription factor Hnf4a in mammalian cardiac development.

## Results/Main

### The scATAC-CoGAPS algorithm

CoGAPS is a sparse, Bayesian matrix factorization algorithm which decomposes a matrix of sequencing data into two output matrices, representing learned latent patterns across all the samples and genomic features of the input data (11), (12). The first of these is called the Amplitude matrix, and it contains a numerical representation of the degree to which each feature contributes to each latent pattern learned by the algorithm. The second is termed the Pattern matrix, which represents the degree to which each learned latent pattern is present in each single cell (Fig. 1A) (15). Latent patterns are intended to capture common accessibility across both genomic features and cells, and thus identify the regulatory biology common among cells in the data (hereafter they will be referred to simply as patterns). The scATAC-CoGAPS algorithm takes as input a count matrix with reads aggregated across any relevant summary feature (e.g. peak regions or DNA motifs that identify TF binding sites).

**Figure 1.**
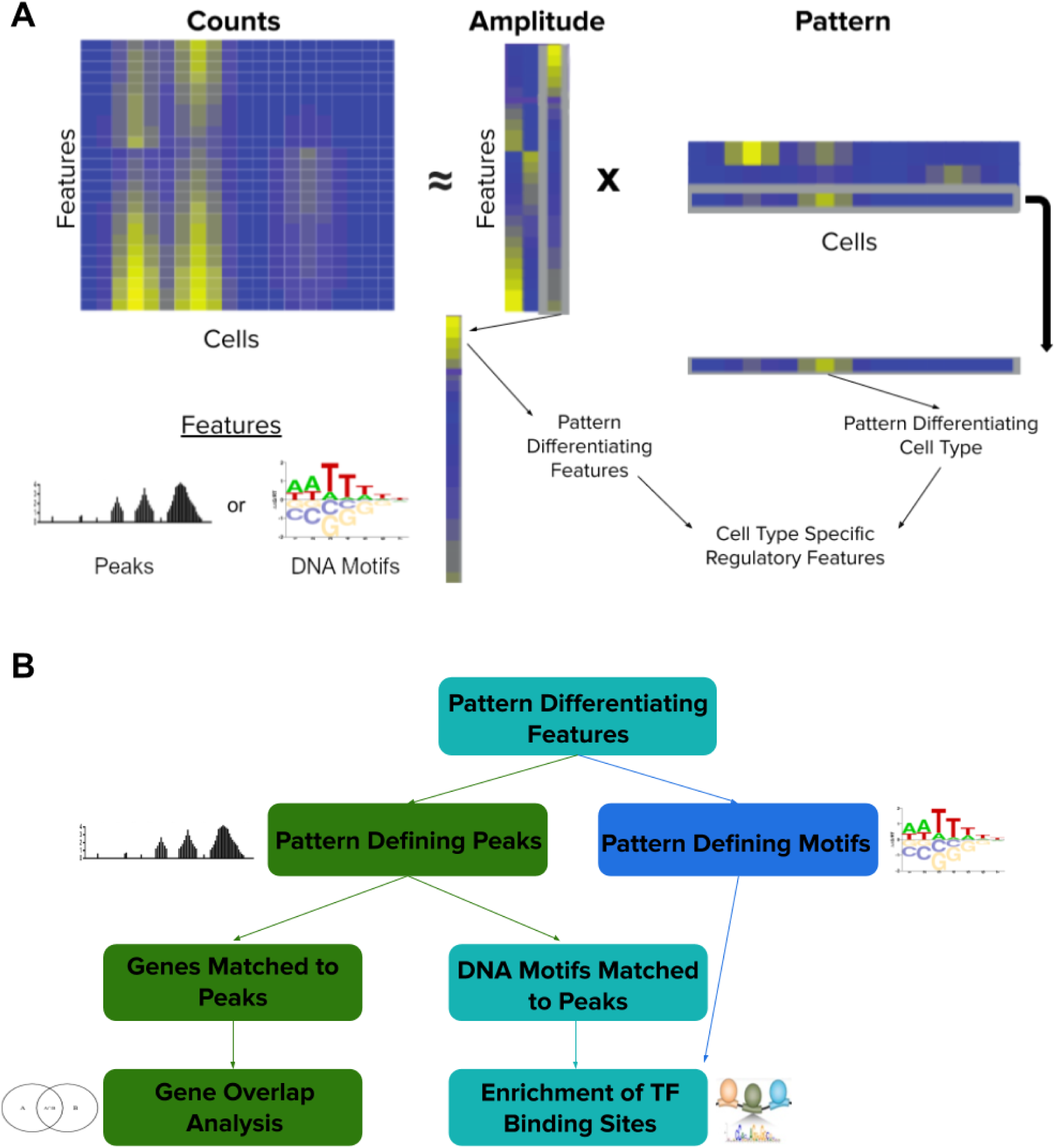
**A** Diagram of Non-negative Matrix Factorization as applied to scATAC-seq data by ATAC-CoGAPS. The Counts matrix (features by cells) is factorized into the Amplitude matrix (features by learned patterns) and the Pattern matrix (learned patterns by cells). The patterns in the Pattern matrix differentiate cell populations, while the same patterns in the Amplitude matrix reveal the differentially accessible features of those cell types. These cell type specific patterns of accessibility can then be used to learn regulatory features that differ across cell populations. **B** Diagram of the analysis approach applied for cell type associated features found by CoGAPS. Features used to produce the input count matrix can be either accessible peaks or DNA motifs. Pattern defining peaks identified by CoGAPS are either matched to genes for gene overlap analysis or matched to DNA motifs to infer TF binding potential. Pattern defining motifs are matched to enriched to TFs, likewise to infer accessible binding sites and thus TF activity in identified cell populations.

The values of the Pattern matrix can be used to distinguish cell types or cell populations specific to each chromatin-accessibility derived pattern. This correspondence allows us to annotate patterns as associated with a particular group of cells. In contrast to standard clustering methods, the patterns learned from CoGAPS can simultaneously identify patterns that delineate individual cell types as well those shared across cell types.

The pattern identified by each row of the Pattern matrix corresponds to a set of gene weights in each column of thee Amplitude matrix. These weights provide information on which specific features (peaks, motifs, etc.) contribute the most to each pattern. In this way, features can be linked to the cell types or cellular states defined by associated patterns, which enables the identification of the active regulatory programs within each group of cells. Further, these learned patterns can be input to our projectR transfer learning method (13), (14) to query their occurrence in related cells in other scATAC-seq datasets.

Assessment of regulatory programs from the amplitude matrix of scATAC-CoGAPS depends upon the features selected for summarization of the scATAC-seq data. The approach outlined here focuses on the annotation of both peaks and DNA motifs. When using open chromatin peaks to define our feature set, we employ two main analysis steps (Fig. 1B). First, we match peaks to genes that fall within the regions they cover, have promoters within these regions, or are in close proximity to these regions. These sets of genes can then be compared to known pathways via gene overlap analysis (16), returning significantly overlapping pathways. Peaks can also be searched for known DNA motifs and their possible TF bindings. The frequency of these potential TF binding sites can inform an understanding of which regulatory effectors are characteristic of a specific cell population. While other analysis methods require one particular mode of feature summarization, CoGAPS allows for the use of any feature that facilitates aggregation of reads into a count matrix. If we instead use a feature space initially defined by DNA motifs, we can again match pattern-defining motifs directly to known TF binding sites to determine enrichment for particular TFs, often extending the number of unique regulatory patterns we are able to uncover from the data (compared to using a peak based feature space alone). However, given that a feature space of peaks provides more options to interrogate regulatory biology (i.e. pathways and TF binding vs TF binding alone), we employ peak summarization as default in our analysis throughout, and utilize a motif-defined feature space to supplement this analysis.

### scATAC-CoGAPS differentiates known cell identities in scATAC-seq data

To demonstrate the capacity of CoGAPS to distinguish cell populations, we run the algorithm on publicly available scATAC-seq data published by Schep et al (6). These data derive from twelve cell cultures, comprising ten different known cell lines (listed in Supplemental Table 1). The cell lines in the data are generally well-characterized, which allows for validation of the cell-type specific regulatory programs predicted by scATAC-CoGAPS. Using peaks to define our feature space, we apply CoGAPS to search for seven patterns of accessibility in the data (see Methods for dimensionality selection). After the factorization, we associate each cell with a single pattern using the PatternMarker statistic included in the CoGAPS package (12). Pattern classifications learned by CoGAPS on this data set align well with *a priori* knowledge of cell line annotations (Fig 2., Supplemental Table 2). Cells belonging to the same cell line are almost always classified within the same pattern (Adjusted Rand Index of 0.90).

**Figure 2.**
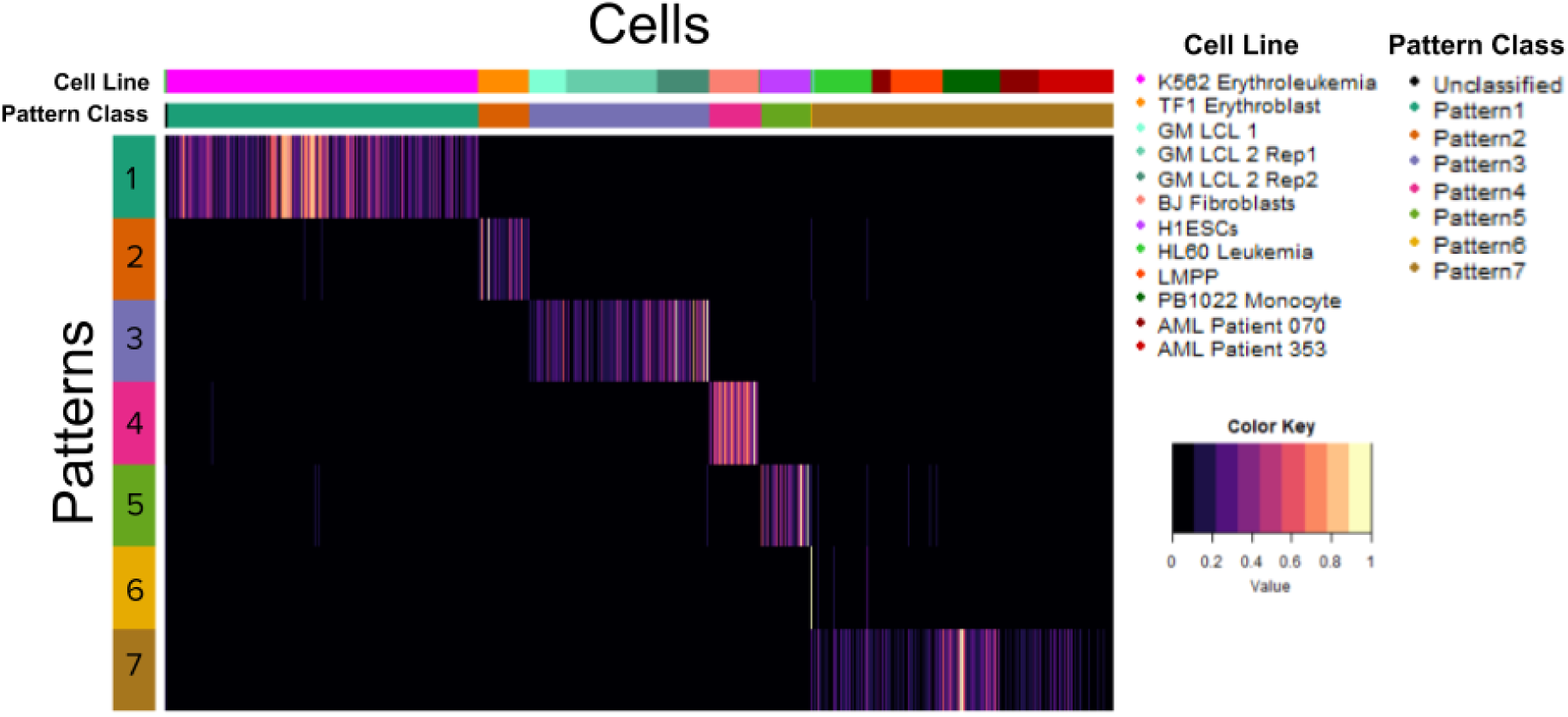
Heatmap of the Pattern matrix with cells matched to learned patterns. The color gradient of the heatmap reflects the Pattern Matrix weights for each cell for each pattern, which indicates the degree to which each pattern is found in each cell, as learned by CoGAPS. Cells are labelled by both Pattern Marker pattern assignment as well as known cell line and culture of origin. Patterns 1-5 all very sharply distinguish a particular cell line. Pattern 6 only captures one cell. Most of the remaining cells are assigned to pattern 7, leaving only 5 cells unclassified.

Pattern 1 and Pattern 2 perfectly classify K562 Erythroleukemia and TF1 Erythroblast cells, respectively. GM B-cell derived LCLs, BJ Fibroblasts, and H1 Embryonic Stem Cells each have 2 or fewer cells misclassified by patterns 3, 4, and 5. We note that Pattern 3 captures all three cultures of GM lymphoblastoid cell lines (GM LCLs), indicating that CoGAPS is differentiating these cell lines via regulatory differences of biology rather than through technical artifacts of cell culture. Pattern 6 is most significantly associated with HL60 Leukemia cells, however, due to the sparse signal in pattern 6, the patternMarker statistic only assigns one HL60 cell to that pattern, and the rest to pattern 7. Pattern 7 is assigned most of the remaining cells in the data, and while it is most significantly associated with PB1022 Monocytes, it also shows signal across HL60 Leukemia cells, Lymphoid-Primed Multipotent Progenitors, and the two AML patient cell lines. We hypothesize that the regulatory similarity derived from the shared hematopoietic origin of these cells is responsible for this common signal.

While the CoGAPS solution described above is for seven patterns, the selection of an optimal dimensionality for unsupervised learning remains an open question, and there probably is no single correct number of patterns to use (17). Therefore, we also run CoGAPS to analyze the scATAC-seq data for additional dimensions. When increasing dimensionality beyond 7, CoGAPS finds patterns that more strongly differentiate Monocytes and Lymphoid-Primed Multipotent Progenitor cells, but still does not return patterns distinguishing the two Acute Myeloid Leukemia patient cell lines apart from Lymphoid Primed Multipotent Progenitors (Supplemental Fig. 1). For example, at the 13-pattern dimensionality, we observe that pattern 1 mainly distinguishes monocytes, while pattern 10 now captures the unifying signal across HL60, LMPP, and AML patient cells. At the same time, with this higher dimensionality, patterns 4, 6, 8, 11, and 13 have very sparse signal and appear to identify only single cells. Thus, we observe a tradeoff at higher dimensions between improved differentiation of cell types and an increased number of sparse patterns. Based on our results across dimensions, we retain the seven-pattern solution for our remaining analyses in order to optimize cell type differentiation while minimizing the number of sparse patterns that are only associated with a few cells.

### Analysis of accessible features predicts regulatory programs consistent with established biology of cell lines

After using CoGAPS patterns from the seven-dimensional solution to define cellular populations, we use the values of the corresponding feature weights in the Amplitude matrix to ascertain which peaks contribute the most to each learned pattern using the PatternMarker statistic. The peaks identified by the PatternMarker statistic reveal the accessible features of the data that themselves strongly distinguish cell types, which we shall refer to as PatternMarker peaks (Fig. 3A). For most cell lines, the accessibility of the PatternMarker peaks learned from CoGAPS analysis better distinguishes the cell lines than the pattern weights themselves. This result suggests that the features CoGAPS learns reflect biologically relevant differences in accessibility between the cell populations that it is stratifying. Due to its increased granularity, this analysis provides further evidence that Pattern 6 is characteristic of HL60 Leukemia cells, and that the peaks associated with Pattern 7 are the most accessible in PB1022 Monocytes.

**Figure 3.**
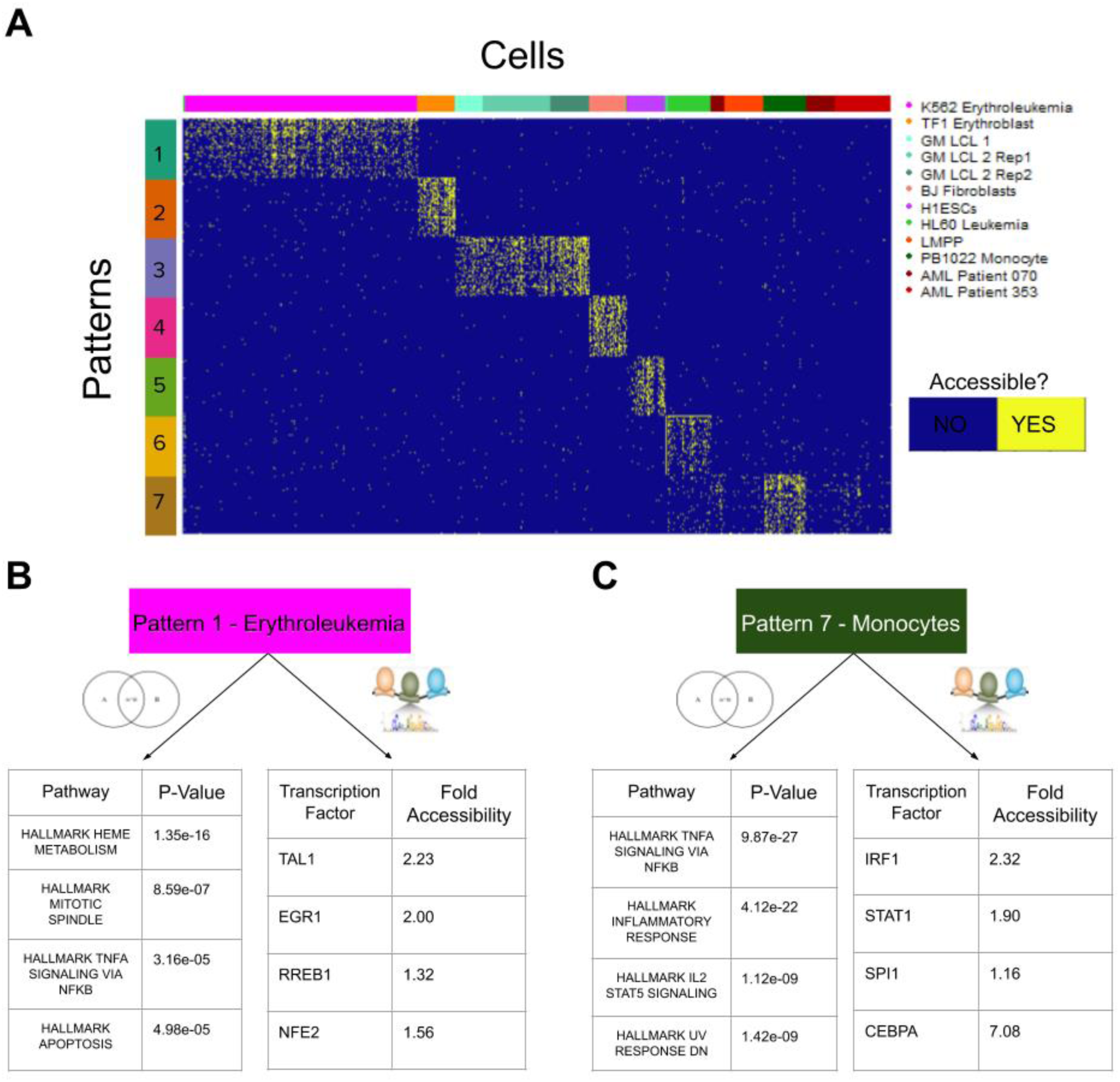
**A** The PatternMarker statistic is used to find the 50 most pattern-distinguishing peaks for each pattern. The counts recorded at these peaks from the scATAC-seq experiment are binarized for accessibility and plotted across all cells in the data. **B**,**C** Examples of the MSigDB Hallmark Pathways with significant overlap to genes matched to PatternMarker peaks (the 4 most significant pathways for each pattern) and Transcription Factors with high numbers of possible binding sites in PatternMarker peaks. TFs listed are those that are both within the top 15 list of TFs with the most enriched binding sites and have highly plausible functional annotations for activity in these cell lines. Fold accessibility refers to the peaks overlapping with the region of the TF gene, relative to other peaks in the K562 Erythroleukemia cell line and PB1022 Monocyte cell line, respectively.

The learned PatternMarker peaks can be associated with cell-specific regulatory mechanisms using pathway and transcription factor enrichment analysis (Supplemental Files 1 and 2). For example, Pattern 1 (the K562 Erythroleukemia-associated pattern) identifies the MSigDB HALLMARK HEME METABOLISM pathway as the most significantly associated with the cell line (Fig. 3B). This matches our biological expectation, as increased accessibility of or near genes associated with Heme metabolism is consistent with the erythroid lineage K562 cells derive from. The second most significant pathway is HALLMARK MITOTIC SPINDLE, which suggests the uncontrolled division of this cancer cell line may be driven by epigenetic changes.

Motif analysis from these accessible peaks further identifies TFs with the most accessible binding sites as potentially active regulators in the pattern-associated cell population. The top 15 TFs enriched within the K562 cell associated pattern include TAL1, EGR1, RREB1, and NFE2 which have all been associated with leukemia (18), (19), (20) or, in the case of NFE2, is an erythroid nuclear factor. TAL1 is a noteworthy hit, as K562 cells were used to establish TAL1 as a driver of leukemia (18), thus providing support for the validity of this approach. To measure the likelihood that the TFs are themselves expressed, we then find the relative accessibility signal at the peaks overlapping the genes of these candidate TFs. All of the above TFs identified from motif analysis also have increased gene accessibility compared to the average peak accessibility in K562 cells, with TAL1 having the highest relative accessibility (Supp. Fig. 2). The accessibility of the gene is most notable for the peak overlapping with the transcriptional start site (TSS) of the gene, with the frequency of the accessibility signal decreasing among the peaks further from the TSS.

The genes overlapping with the peaks that contribute most strongly to the Monocyte-associated Pattern 7 are enriched for the MSigDB HALLMARK INFLAMMATORY RESPONSE and HALLMARK TNFA SIGNALING VIA NFKB pathways (Fig. 3C). Both pathways are biologically consistent with the known role of monocytes in immunity and inflammation, as well as with the immunological roles of the other hematopoietic lineage cells secondarily associated with Pattern 7. Within the top 15 TFs with the most enriched binding sites, IRF1, STAT1, CEBPA, and SPI1 all have previously established roles in the regulation of monocytes (21), (22), (23), (24) and all TF genes have increased gene accessibility relative to average for monocyte peaks in the data (Fig. 3C). The pathway and TF enrichment results for all other patterns are listed in Supplemental Files 1 and 2. Taken together, these results demonstrate the capacity of scATAC-CoGAPS to identify regulatory features of biological relevance from scATAC-seq data.

### Summarization of the count matrix by DNA motifs extends the regulatory patterns CoGAPS learns from scATAC-seq

While using peaks as summarization of ATAC-seq reads provides more avenues for downstream analysis, it has been previously shown that motif-level summarization is an additional information rich feature space for scATAC-seq analysis (6). Therefore, we compare our previous peak-level CoGAPS analyses for the Schep et al. data set (6) to motif-based CoGAPS analyses (labeled Pattern Defining Motifs in Fig 1B) of the same dataset to assess the impact of feature selection on the inferred regulatory programs. CoGAPS analysis of this motif-based count matrix identified 10 total patterns from the data (Supplemental Figure 3A). Patterns 4, 6, and 8 from this motif-level CoGAPS run differentiate GM-LCLs, BJ Fibroblasts, and TF1 Erythroblasts, respectively.

The other patterns identify additional cell populations that are not found when the data are analyzed using peak feature space (Supplemental Fig. 3A). For example, Pattern 10 identifies regulatory similarity between K562 Erythroleukemia cells and TF1 Erythroblasts, a pattern that peak based analysis does not find (Supplemental Fig. 3B). In Pattern 10, we identify high enrichment of candidate TF binding sites for GATA transcription factors, which are known to have critical roles in erythroid differentiation and are shared between Erythroleukemia and Erythroblasts (25). We additionally find that the PatternMarker motifs identified by CoGAPS in this analysis are nearly all different than the motifs found by peak-based analysis. When patterns that seem to differentiate the same cell types are compared, less than 10% of the motifs identified by each analysis overlap (overlap for Fibroblast associated patterns is given in Supplemental File 3).

These results suggest that using DNA motif-based summarizations identifies additional regulatory information from the same cell types contained within the same data, and directly supports the use of both peak and motif based summarizations to fully characterize the regulatory biology of cellular subpopulations in scATAC-seq data. Notably, motif-based summarization appears to better identify patterns of accessibility that are shared across multiple cell types, while peak-based summarization better differentiates individual cell types.

### Transfer Learning with projectR establishes the generality of the regulatory programs CoGAPS patterns capture

Once we have established signatures of accessibility for cell populations in our data, we employ transfer learning with the R/Bioconductor package projectR (13), (14) to determine whether these signatures appear in similar cell populations from other experiments. Notably projectR can efficiently detect the presence of previously learned patterns of accessibility in separate scATAC-seq data as a means of *in silico* validation and discovery. This capability allows for the development of cell population-specific accessibility signatures based on CoGAPS results, which can be used to test for regulatory programs of interest in novel samples.

We demonstrate projectR’s application to scATAC-seq by transferring the patterns learned in peak-level summaries of the Schep et al. (6) cell line data to scATAC-seq data from Buenrostro et al. (26), which contains 10 different hematopoietic lineage cell types labelled via Fluorescence Activated Cell Sorting (Supplemental Table 3). We project the monocyte-associated pattern (Pattern 7) from the Schep et al. data onto the Buenrostro et al. data and observe that the monocytes in the target data are most significantly associated to the accessibility pattern (Fig. 4A). Comparing average cell line association with the pattern in the target data may make the specificity of the monocyte association more visually clear (Supp. Fig. 4). As previously noted, there is considerable Pattern 7 signal among other non-monocyte hematopoietic-lineage cells within the Schep et al. data set, and this is reflected in the general signal observed in the Buenrostro target data set.

**Figure 4.**
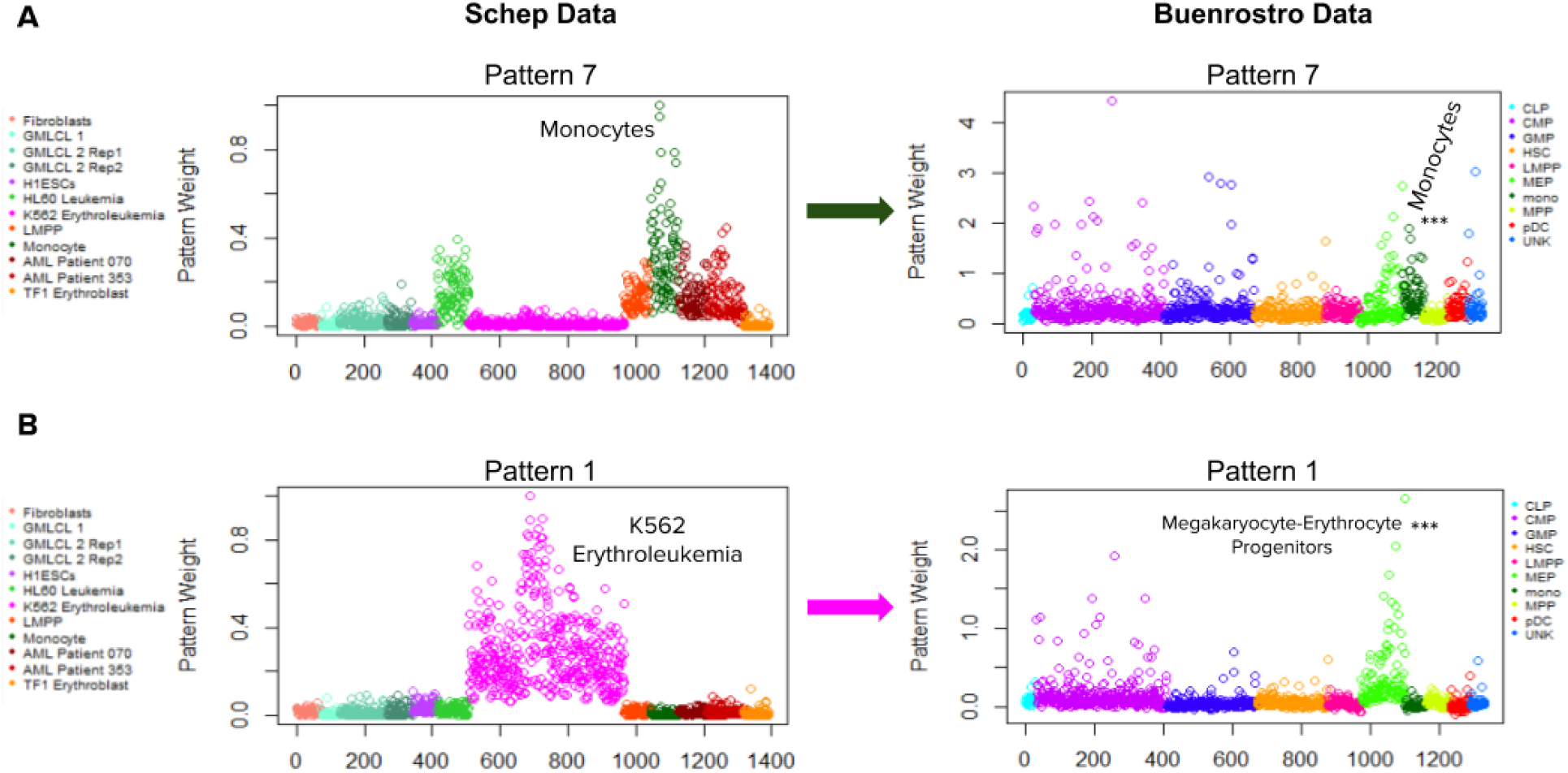
**A** Projection of peak accessibility associated primarily with monocytes in the Schep data set into the Hematopoietic lineage Buenrostro data set. The Monocytes in the Buenrostro set are the cell type most significantly associated with the pattern, as determined by a pairwise Wilcoxon Rank Sum Test. **B** Projection of the accessibility signature associated with the K562 Erythroleukemia cell line in the Schep data into the hematopoietic lineage data. This signature is most significantly associated with Megakaryocyte-Erythrocyte Progenitor cells.

ProjectR can also provide information on the regulatory overlap between different cell types. In this case, it provides insight into the regulatory similarity between two distinct cell populations. For example, projection of the K562 Erythroleukemia cell line pattern from the Schep et al. data (Pattern 1) into the Buenrostro et al. data has the strongest signal in Megakaryocyte-Erythrocyte progenitors (Fig. 4B). This observation supports the presence of overlapping patterns of accessibility between these two populations, consistent with the expected regulatory similarity between Erythroleukemia and Erythrocyte progenitor cells.

### Analysis of matched scRNA-seq data validates regulatory programs learned from scATAC-CoGAPS

When scRNA-seq data is available for cells from the same experimental conditions as scATAC-seq data, we can validate ATAC-CoGAPS predicted TF activity using transcription data of known TF gene targets. CoGAPS can be applied to the matched scRNA-seq data to find pattern-defining genes for each cell population as described in (12). These genes can be ranked on the basis of their contribution to each pattern (using the PatternMarker statistic), and then tested for enrichment in the set of genes known to be regulated by a candidate TF using Gene Set Enrichment Analysis (GSEA) (27) (Supp. Fig. 5). In this analysis method, genes known to be regulated by a TF are used as the “pathways” input for GSEA with the ranked PatternMarker genes.

No matching scRNA-seq data was available for the Schep et al. data set. Therefore, we sought to validate this method using matched scRNA-seq and scATAC-seq data from mouse embryonic cardiac progenitor cells at days 8.5 and 9.5 of development, as described by Jia et al. (28). We run CoGAPS on both data sets to learn 7 patterns in peak-level summarized scATAC-seq data and 6 patterns in the scRNA-seq data. There is much more regulatory similarity than dissimilarity between cardiac progenitors only one day apart in development, and thus the most distinctive patterns we find in the scATAC-seq data set are those that reflect sustained open chromatin across days 8.5 and 9.5 of development (Patterns 1 and 7) (Supp. Fig. 6). As patterns 3 and 6 from the scRNA-seq experiment also have signal across all cells in the data, we continue by comparing the patterns found across cells rather than the patterns that stratify distinct cell populations. To make this comparison, we first find TFs enriched within the scATACseq data for all cells, and then list the genes known to be regulated by each of the TFs. Then, we find the PatternMarker genes from scRNA-seq from the patterns that show signal across all cell types (patterns 3 and 6). GSEA between the sets of genes regulated by the predicted TFs and the PatternMarker genes provides significant support for Tbx20 TF activity (FDR adjusted p-value of 0.015) and Hnf4a activity (FDR adjusted p-value of 0.042) across these developing cardiac cells (Fig. 5A, 5B). Tbx20 plays a major role in cardiac development (29), which is consistent with the known biology of embryonic cardiac cells. A homologue of Hnf4a was recently shown to play an important role in normal embryonic development of the chicken heart (30). This result corroborates that finding and suggests that Hnf4a may play a role in cardiac development across a wide phylogenetic range; particularly that it acts in mammals as well.

**Figure 5.**
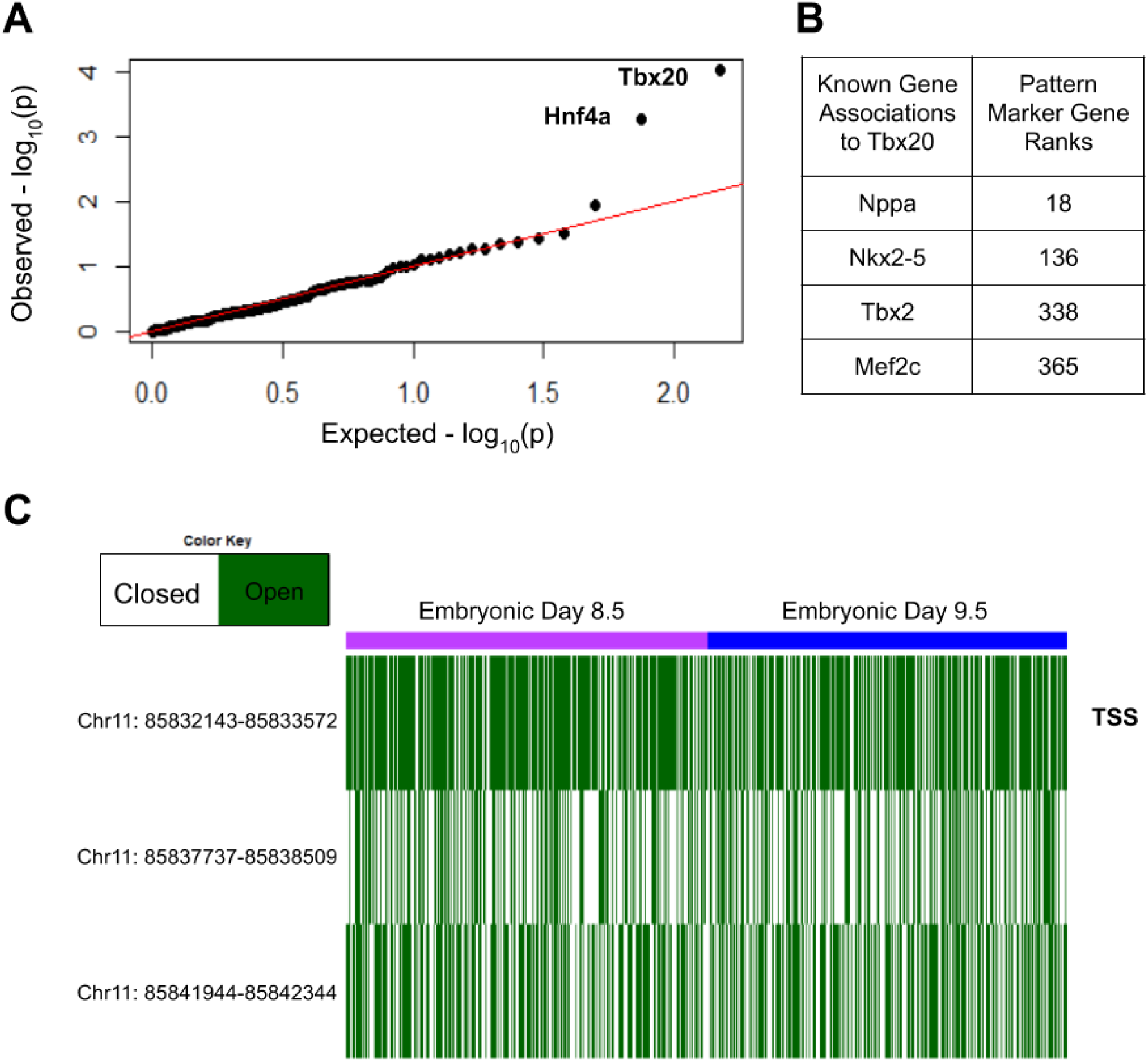
**A** qqPlot of p-values for gene set enrichment analysis of the Transcription Factors’ gene networks predicted from scATAC-seq CoGAPS and the genes ranked by scRNA-seq CoGAPS. **B** Known genes regulated by Tbx20 and their PatternMarker ranks from CoGAPS analysis in matched scRNA-seq. **C** Accessibility at the Tbx2 gene in the scATAC-seq data, showing the correspondence of its accessibility and expression levels across mouse cardiac progenitor cells, at embryonic days 8.5 and 9.5. The Transcriptional Start Site overlapping peak (marked with TSS) is the most consistently accessible.

To investigate the accessibility of genes associated with Tbx20 using scRNA-seq, we find overlapping peaks of said genes within matched scATAC-seq data. The peaks corresponding to the Tbx2 gene and the Nkx2-5 gene are accessible across the cells in the data (fold accessibility 2.39 and 1.51, respectively), while Mef2c and Nppa peaks are less accessible than average (fold accessibility 0.84 and 0.30) (Fig. 5C, Supp. Fig. 7). The Tbx2 gene is particularly accessible in the peak overlapping with its transcriptional start site (fold accessibility 3.11). The lack of accessibility among the Mef2c and Nppa genes suggests that accessibility and gene expression do not always align, though we do observe general correspondence between the two data modalities, particularly in transcriptional start site overlapping peaks.

## Discussion

Single-cell epigenomics methods such as scATAC-seq capture a wide array of regulatory features genome wide, but our ability to extract this information is still limited. Here we present the application of CoGAPS and projectR to scATAC-seq, providing an analysis framework for Bayesian Non-Negative Matrix Factorization to uncover regulatory information from sparse, high-dimensional epigenomics data and project these learned patterns across data sets and sequencing platforms.

CoGAPS (Coordinated Gene Expression in Pattern Sets) was originally developed for the analysis of gene expression data. The ability of CoGAPS to extract relevant patterns from different data sources is a great strength of the algorithm. Here, we leverage this capacity to develop a basic framework for integrative analysis of multiple scATAC-seq and scRNA-seq data sets. Since CoGAPS can be applied to any sequencing technology that can produce a count matrix, this framework we present has the potential to support the integrated analysis of additional multi-omics data sets. The importance of this capacity continues to grow with the increasing affordability and concomitant ubiquity of sequencing technologies, and the massive and varied data sets such technologies produce. Furthermore, CoGAPS allows for the summarization of reads to any relevant genomic feature (e.g. peaks, DNA motifs, etc.) and facilitates the learning of a wider range of regulatory patterns than methods that require a specific summarization method.

This study presents CoGAPS and projectR as a paired set of tools for cross-study analyses of regulatory biology from scATAC-seq data. The projectR transfer learning software is broadly applicable for features learned with unsupervised methods in addition to CoGAPS (14). This flexibility of projectR will support further cross-study analyses with emerging scATAC-seq methods (15). While this study demonstrates the robustness of CoGAPS for inferring regulatory biology from scATAC-seq data, we resolve different aspects of that biology at different dimensionalities and data summarizations. We hypothesize that accounting for these features across hyperparameters as well as additional features informed from ensembles of features learned from alternative methods are critical to resolve the complex landscape of regulatory biology encoded in the data, consistent with emerging literature on multi-resolution methods (31).

We find that TF motif-based analysis tends to find more patterns that have signal across cell types, while peak-based analysis finds more cell type specific signal. We hypothesize that each peak mostly contains signal corresponding to one or a few genes, and therefore peaks more finely map cell populations to distinct cell types. Transcription factor motifs, on the other hand, contain signal corresponding to larger regulatory changes that are more likely to be shared between cell types, and thus analysis in this space yields more patterns with signal across cell types. If this hypothesis is correct, it seems possible that an enhancer-based space could provide another higher order feature, that could identify more patterns of regulatory biology that act across multiple cell types.

The projectR software package makes it possible to determine whether the patterns learned in one data set are present in others, and can do so in a way that is fast and easy to implement. This a major strength of the approach we present, as it helps to simultaneously extend and validate learned regulatory patterns, while also allowing for the comparison of regulatory biology in multiple scATAC-seq data sets. Most current scATAC-seq analysis methods are limited in application to a single data set and any results cannot be directly related to other data sets or analyses. This fact severely limits the efficiency of broad analyses, and the information that can be learned from distinct but complementary data sets. ProjectR thus synergizes with CoGAPS and has tremendous potential for use in analyzing disease-specific data sets. For example, if we can establish robust signatures of disease or treatment associated biology, such as genomic dysregulation and markers of drug efficacy, respectively, we can use CoGAPS and projectR to leverage clinical data for an improved understanding of disease mechanisms (32), (33) and to guide treatment decisions.

Matrix factorization is well suited to the problem of understanding scATAC-seq data, as the technique learns patterns that distinguish both features and cells within the two factorized output matrices. This output is conducive to a more thorough analysis of the regulatory differences between the cell populations in the data than most available methods can provide. Thus, it is unsurprising that matrix factorization has been previously applied to scATAC-seq analysis (34), (35), (36). We use CoGAPS because the Bayesian optimization of the factorization has been previously shown to be more robust to initialization than gradient-based NMF, resulting in more biologically relevant patterns (11), (12), (37). Duren et al. and Zeng et al. each apply a coupled factorization for integrative analysis of multiple sequencing modalities, allowing for simultaneous clustering and investigation of regulatory biology (34), (35), (36). ProjectR can potentially be applied to the output of these coupled factorizations, allowing for transfer of these integrated patterns of regulatory biology across data sets. Coupled factorization may be a promising avenue for future development of integrative analysis with CoGAPS, and projectR will be able to serve in this context to determine whether different coupled factorization methods identify similar patterns of regulatory biology.

We note that multi-platform data integration is a broad area of research, extending well beyond matrix factorization based approaches. Coupled correlation analysis has recently been applied to scATAC-seq and scRNA-seq, both allowing for integrative analysis and imputation of spatial transcriptomics information (38). Linked Self-Organizing Maps have also been used in this context, providing the capacity to find differences between relatively similar cell types (39). In the area of experimental methods development, recent research has provided techniques for parallel sequencing of RNA, accessibility, and methylation from single cells, vastly lowering both the time and monetary cost of joint profiling of single cells (40), (41). Further, multiple efforts are underway to sequence transcriptomics and chromatin accessibility from the same single cell, which promises to improve the fidelity of multimodal analysis and the ability of multi-omics computational methods to learn the regulatory biology of constituent cell populations.

## Conclusions

The ATAC-CoGAPS analysis framework provides robust tools for identifying regulatory biology from scATAC-seq data. Further, it provides the capacity for integrative multi-omics analysis, as well as Transfer Learning of accessibility signatures across data sets. These characteristics allow the ATAC-CoGAPS framework to produce consensus signatures of cell populations that apply across sequencing modalities and across variations in cellular conditions, which is infeasible with other currently available methods.

## Methods

### ATAC-CoGAPS Pipeline

The ATAC-CoGAPS software is freely available as an R package from https://github.com/FertigLab/ATACCoGAPS. Briefly, this software package includes functions for preprocessing of scATAC-seq data to run the CoGAPS algorithm (version ≥ 3.5.13), as well as functions for subsequent analysis of the results. Each of the steps taken to perform the standard ATAC-CoGAPS workflow are demonstrated at https://rossinerbe.github.io/. All analyses performed to produce the results described in this work are available from https://github.com/rossinerbe/ATACCoGAPS-Analysis-Code.

Input reads from a scATAC-seq experiment are summarized into some feature space (peaks, DNA motifs, etc.) and into an input count matrix, features by cells. Specific preprocessing steps are outlined in the analysis code linked above. Next, the count matrix is input to the R/Bioconductor package CoGAPS. CoGAPS employs a sparse, Bayesian non-negative matrix factorization algorithm to decompose the scATAC-seq count matrix **C**, features by cells, into an Amplitude matrix **A**, features by learned patterns, and a Pattern matrix **P**, learned patterns by cells as described in (11) and (12). The primary parameter for the application of CoGAPS is then the feature level summarization and number of learned patterns, described in further detail below. To account for sparsity, we filter this input count matrix **C** is filtered to remove any feature or cell that is more than 99% zero.

The next steps of the ATAC-CoGAPS analysis framework then focuses on the output **A** and **P** matrices. Unless otherwise noted, all steps are functionalized within the ATACCoGAPS package and all outside packages used are wrapped within ATACCoGAPS functions (see the workflow at https://rossinerbe.github.io/ for detailed implementation with code). We first evaluate the results object from CoGAPS by plotting the Pattern matrix **P** (learned patterns by cells) to determine which patterns differentiate which cell populations. Annotations of patterns to cell populations are made using the PatternMarker statistic to determine the pattern each cell is most defined by, thereby clustering cells to each pattern. Alternatively, if *a priori* determined cell populations are known (e.g. by fluorescence activated cell sorting) we can determine which of these populations have significant signal in a pattern by calling the pairwise.wilcox.test R function for each pattern (not functionalized in ATAC-CoGAPS). The Adjusted Rand Index is used to quantify the overall clustering of CoGAPS on the Schep et al. data set (6) using the pattern to cell line annotations listed in Supplemental Table 2. Once these correspondences of pattern to cell type are annotated, we can then turn to the Amplitude matrix **A** (features by learned patterns). We apply the PatternMarker statistic to find the accessible features that most strongly contribute to each pattern, and thus most define the cell population they distinguish. The number of features used in these analyses is determined by thresholding of the PatternMarker statistic such that the feature is assigned to the pattern for which its association is scored most highly (12).

Analysis of the amplitude matrix **A** also depends critically on functional annotation. If peaks are used as summarization, we first match peaks to genes within or near those regions using the GenomicRanges R package version 1.36.1 (42). We then find enrichment of those genes within known pathways from MSigDB (in this work we demonstrate this capability using Hallmark Pathways v7.0) (27), (43) using the GeneOverlap R package version 1.20.0 (16).

Additionally, peaks are matched to DNA motifs with potential TF binding sites using the motifmatchR Package version 1.6.0 (6). TFs with common possible binding sites in multiple PatternMarker accessible regions are returned, along with functional annotations, so the biological plausibility of a TF’s activity in a particular cell population based on known function can be considered alongside the enrichment results. Next, the accessibility of the peaks overlapping with the TF gene itself is evaluated relative to the general accessibility of peaks for that cell population to provide evidence as to whether the TF itself is expressed. For each peak that overlaps with the TF gene, the number of cells with accessible reads are counted within the cell population of interest. This number is averaged for all peaks overlapping the TF gene and then this average is divided by the average quantity of accessible cells for all peaks in the cell population. The resultant fold accessibility value is not intended as a precise quantification, but rather an approximate guide to assess whether a gene is generally accessible in a particular cell population.

If the data is summarized to motifs before running CoGAPS using ATACCoGAPS preprocessing functions (which employ motifmatchR for motif matching), the downstream analysis is performed similar to the above. Common TF bindings are returned and assessed for relative accessibility to determine whether the TFs are likely to be themselves expressed in the cell population. Relative accessibility of the TF genes is calculated as described previously.

Learned patterns can be projected into other data sets to determine if the signatures identifying cell populations within one data set apply more generally. We use the projectR package version 1.0.0 (13), (14) to perform this analysis. If we use a peak feature space for transfer learning, peaks in the target data set must be matched to peaks in the source data set to project the patterns learned in the source data set. We use the set of all peaks that have any overlap between the two sets as the features we project from and into. If we instead apply DNA motifs as the feature space, all motifs that occur in both data sets are used for projection.

We apply CoGAPS to scRNA-seq data in order to validate candidate TFs identified by scATAC-seq analysis. First, patterns that distinguish the same cell populations are identified. Then, the PatternMarker statistic is used to rank the scRNA-seq genes most associated to each pattern. The TFs identified as described above in scATAC-seq are matched to annotations from the TRRUST database version 2 (44) which list the genes the TFs are known to regulate. These gene sets are compared to the scRNA-seq CoGAPS based gene rankings by gene set enrichment analysis implemented with the fgsea R package version 1.10.1 (45). TFs with significant enrichment of the genes they are known to regulate are considered to be supported by multimodal analysis.

### CoGAPS Hyperparameters

All CoGAPS analyses presented in this manuscript are performed with CoGAPS version 3.70.0. Factorizations are performed in parallel across random subsets of features using the genome-wide option (12) (which should be used unless there are more cells the features, in which case the single-cell option should be used instead) and 10,000 iterations. The only remaining free input parameter for CoGAPS is then the number of patterns, **n**, to learn from the data. The input matrix is features by cells, the Amplitude matrix is features by **n**, and the Pattern matrix is **n** by cells. We note that selecting the number of patterns for unsupervised learning methods is an open question in machine learning. Previously, we and others have found that pattern robustness and chi-squared statistics for goodness of fit across a range of values of **n** provide performance metrics for selection of **n** (46), (47). *A priori* knowledge of the set of conditions or populations each cell derives from can provide an initial heuristic for the selection of **n**. Several CoGAPS runs can be performed in parallel to test different numbers of patterns. After these CoGAPS runs, a Chi-squared test can be performed on the output to determine the goodness of fit of the results and provide numerical guidance on the question of how well different numbers of patterns fit the data.

#### Public Data

This study presents analyses on publicly available scATAC-seq data from (6) (GSE99172), (26) (GSE96769), and (28) (https://github.com/loosolab/cardiac-progenitors on 8/7/2019). In all cases, data were obtained at peak summary (see papers for alignment and peak calling details). Both the Schep et al. 2017 and the Buenrostro et al. 2018 scATAC-seq datasets were downloaded with peaks of equal width. The peaks called for the Jia et al. 2018 data set were not of equal width, so counts were normalized by dividing the values of each peak by its nucleotide width. Motif counts were obtained using ATACCoGAPS software to convert peak counts to motif counts. The scRNA-seq data set from Jia et al. 2018 contains matched single cells to the scATAC-seq dataset. These data were also obtained from https://github.com/loosolab/cardiac-progenitors on 8/7/2019 as normalized counts. Prior to running CoGAPS, all peaks and cells that were more than 99% sparse were filtered out of the data (32,789 peaks and 528 cells for the Schep et al. data set and none for the Jia et al. data set (as it was pre-filtered by Jia et al.)). CoGAPS was run for 7, 13, and 18 patterns in this work on the Schep et al. 2017 data set. CoGAPS was run for 7 patterns on the scATAC-seq data set and 6 patterns on the scRNA-seq data from Jia et al. 2018.

## Supplemental Figures

**Supplemental Figure 1.**
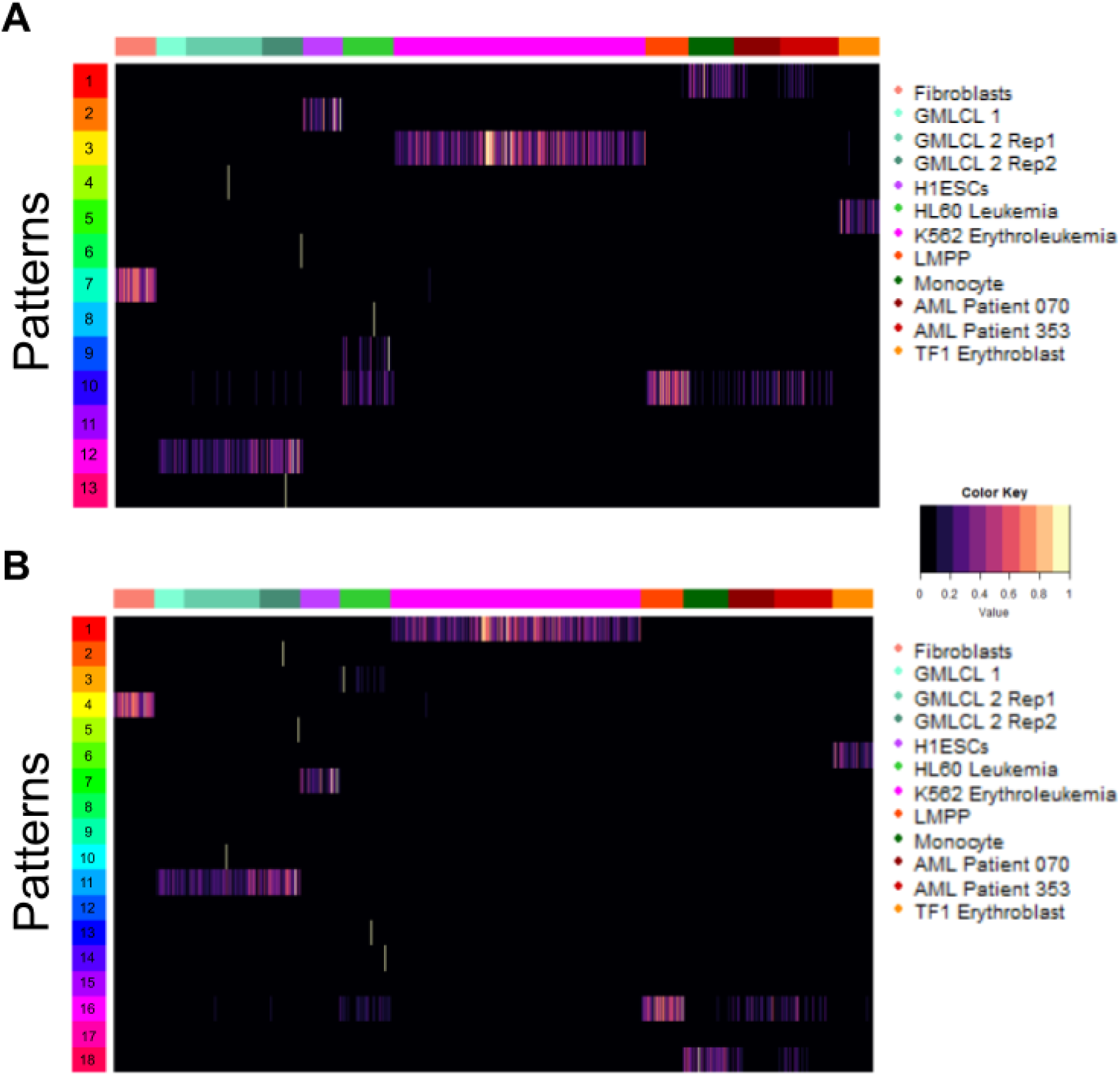
The Pattern matrix is plotted for CoGAPS runs using **A** 13 and **B** 18 patterns for the Schep et al data. Several sparse signal patterns are observed. The monocyte pattern becomes more clear and a LMPP and patient leukemia pattern emerges, which are not seen when running the algorithm for 7 patterns. Pathway enrichment and TF prediction results are robust across different pattern numbers (ie. the patterns that distinguish the same cell types return the same most significant pathways and most enriched TFs for patterns defining the same cell lines) (see Analysis Code).

**Supplemental Figure 2.**
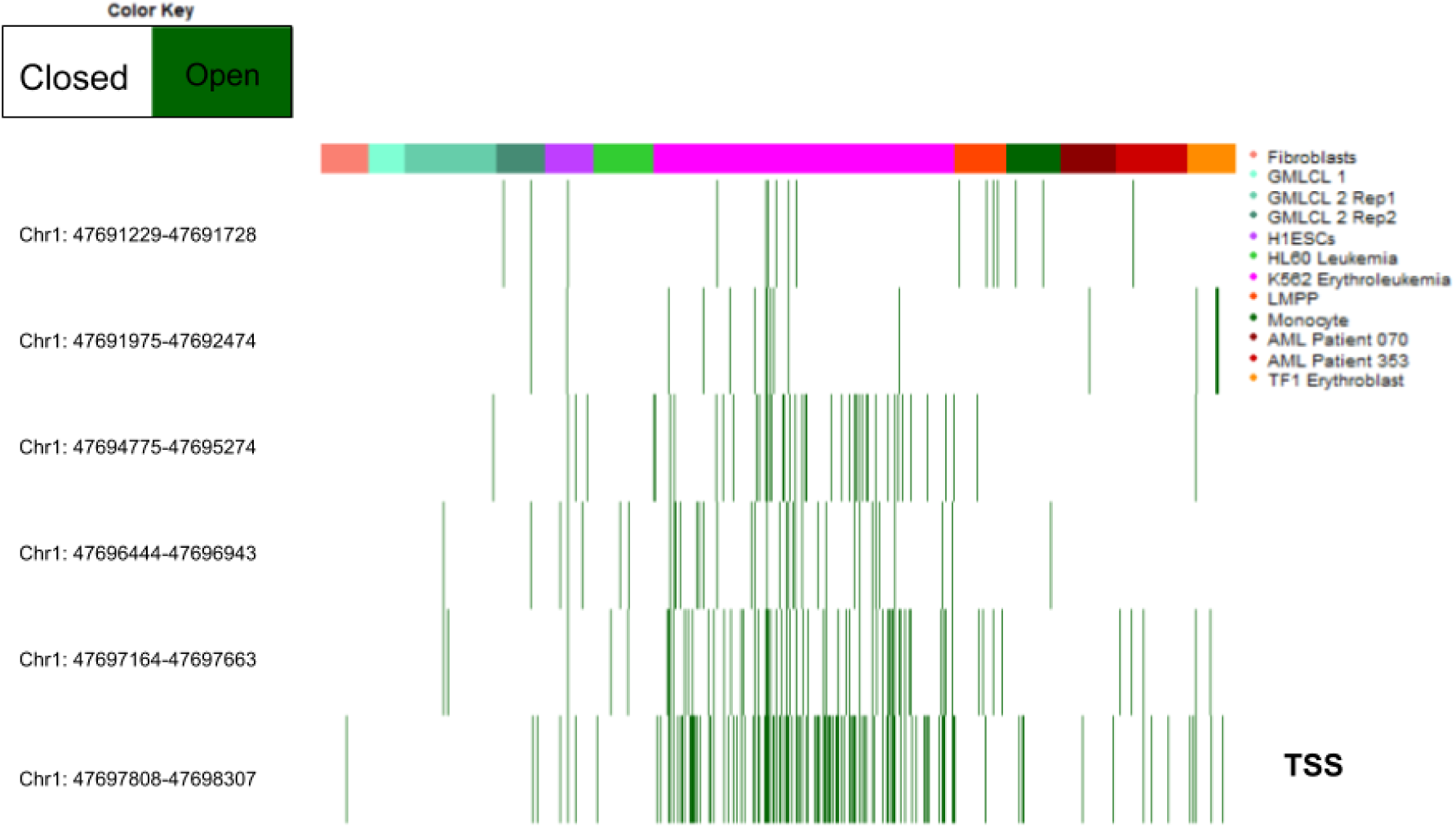
Plot of the binary accessibility of TAL1 overlapping peaks, revealing higher accessibility in K562 Erythroleukemia cells and providing evidence of its specific expression in that cell line. The peak overlapping with the Transcriptional Start Site is marked as TSS and is more consistently accessible among K562 cells than any other TAL1 overlapping peak.

**Supplemental Figure 3.**
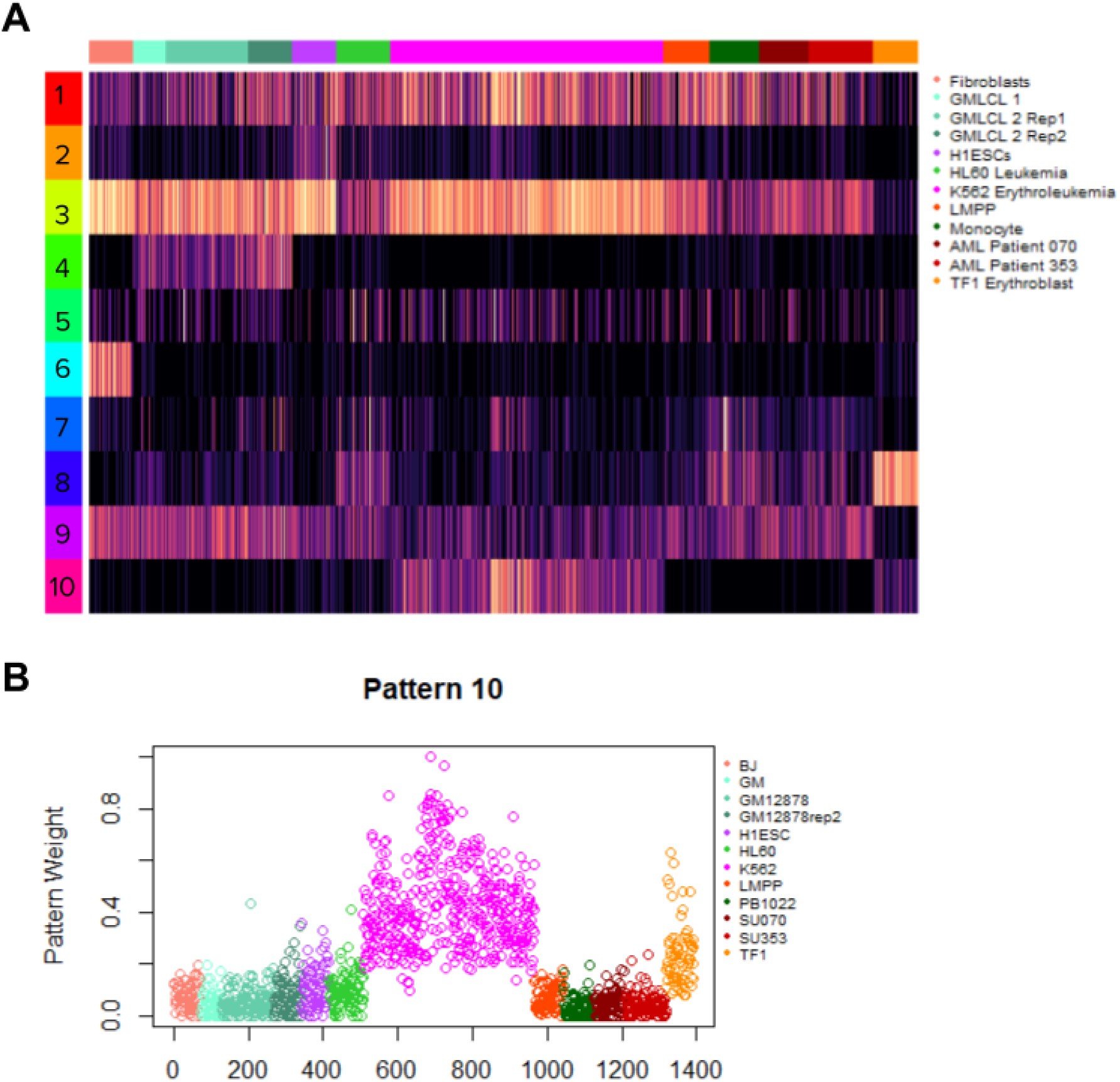
**A** Plot of the Pattern matrix after running CoGAPS with DNA motif summarization on the Schep et al data. The only parameter differences from the peak summarization are that this data is run for 10 patterns and it is run across fewer parallel cores due to there being fewer motifs than peaks. **B** Plot of a pattern found by CoGAPS in the Schep data set when it was run using motif summarization rather than peak summarization (the same as the 10th pattern plotted in **A**, plotted alone for increased visual clarity). Both TF1 erythroblasts and K562 Erythroleukemia cells are strongly associated with this pattern. We do not identify a similar pattern with summarization to peaks.

**Supplemental Figure 4.**
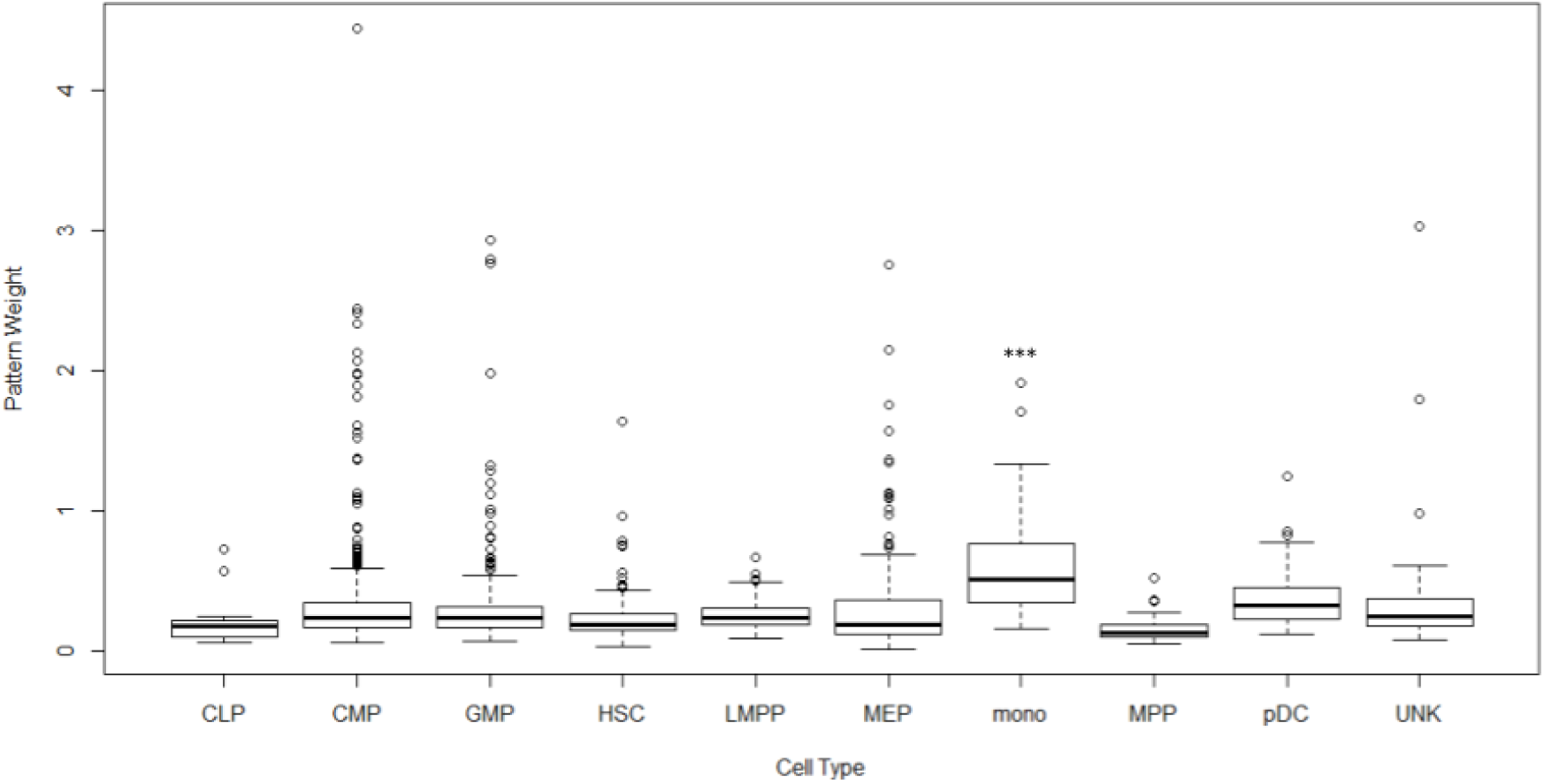
Boxplot of the pattern weight for the transfer of the monocyte associated pattern from the Schep data into the Buenrostro data. The monocytes in the Buenrostro data are most significantly associated with the pattern as evaluated by a Wilcoxon Rank Sum Test.

**Supplemental Figure 5.**
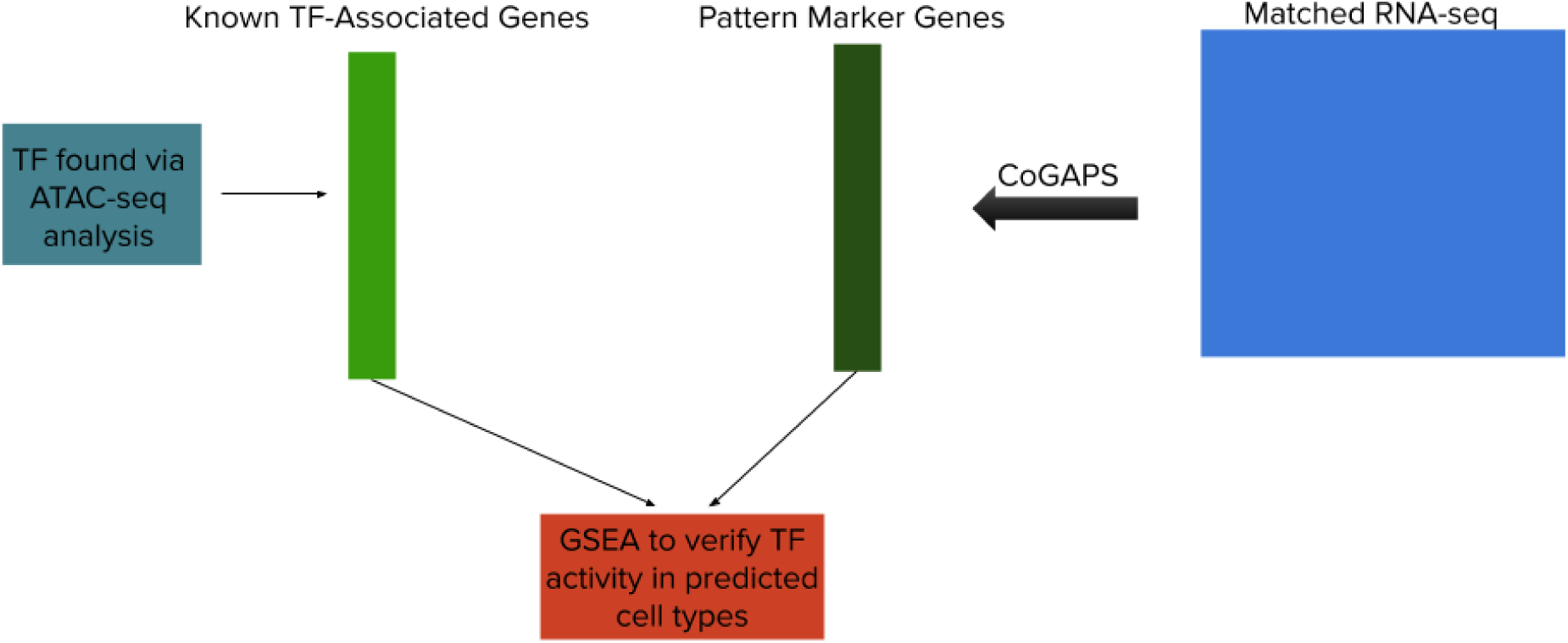
Diagram of the analysis method employed to validate ATAC-CoGAPS candidate using matched scRNA-seq data. The TFs identified by ATAC-CoGAPS are first matched to the sets of genes they regulate. Then, CoGAPS is run on the scRNA-seq data and PatternMarker genes are identified. GSEA is performed between the TF gene sets and the PatternMarker genes to provide transcription-based validation of TF activity.

**Supplemental Figure 6.**
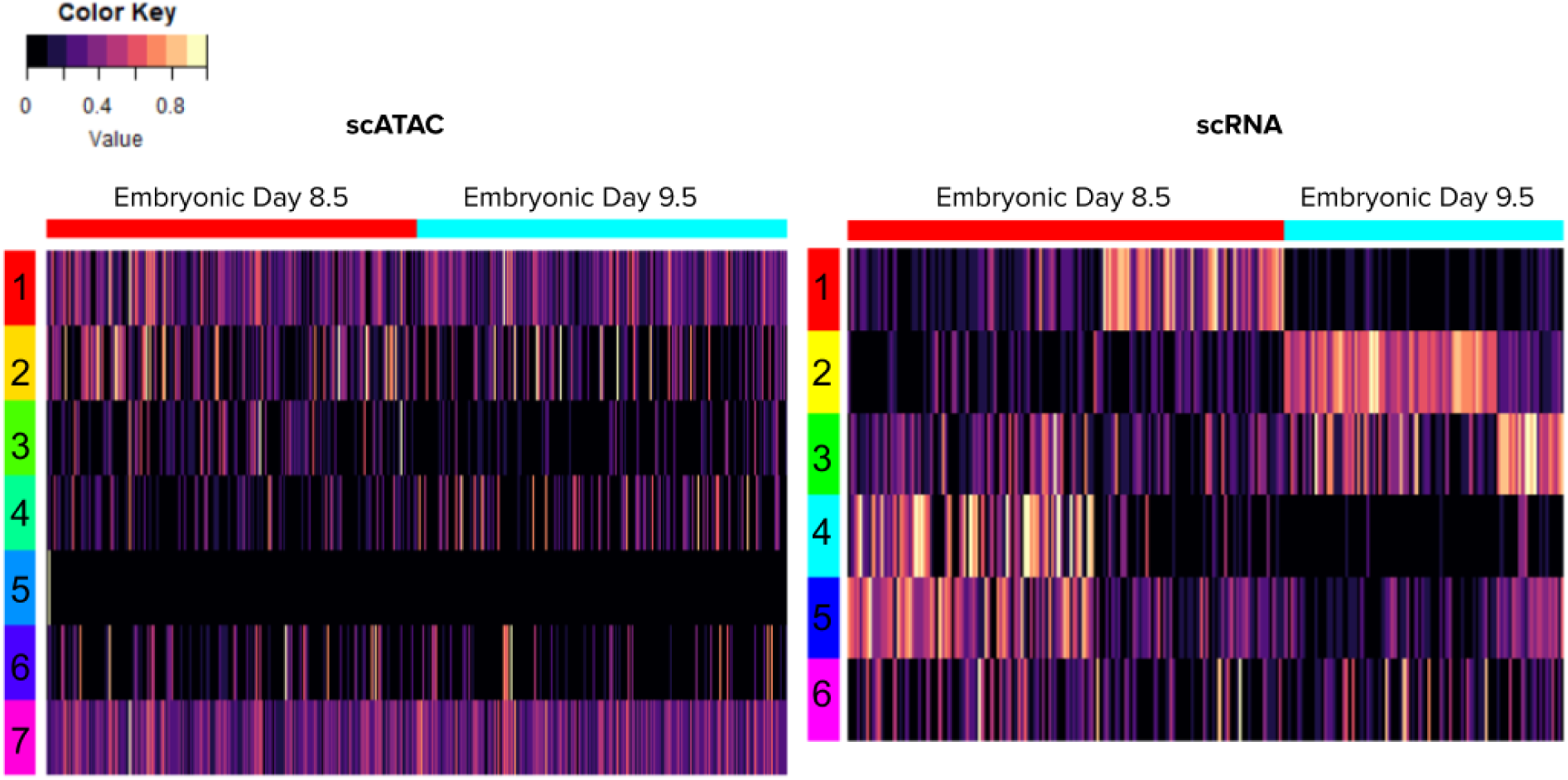
The Pattern matrices plotted for both scRNA-seq and scATAC-seq from matched cardiac development data derived from mouse embryos and published by Jia et al. scRNA CoGAPS finds more differentiating patterns, while most of the scATAC patterns are unifying across the similar cell types, suggesting scRNA-seq is either identifying populations subtypes that scATAC does not capture or is identifying batch effects in the RNA-seq data.

**Supplemental Figure 7.**
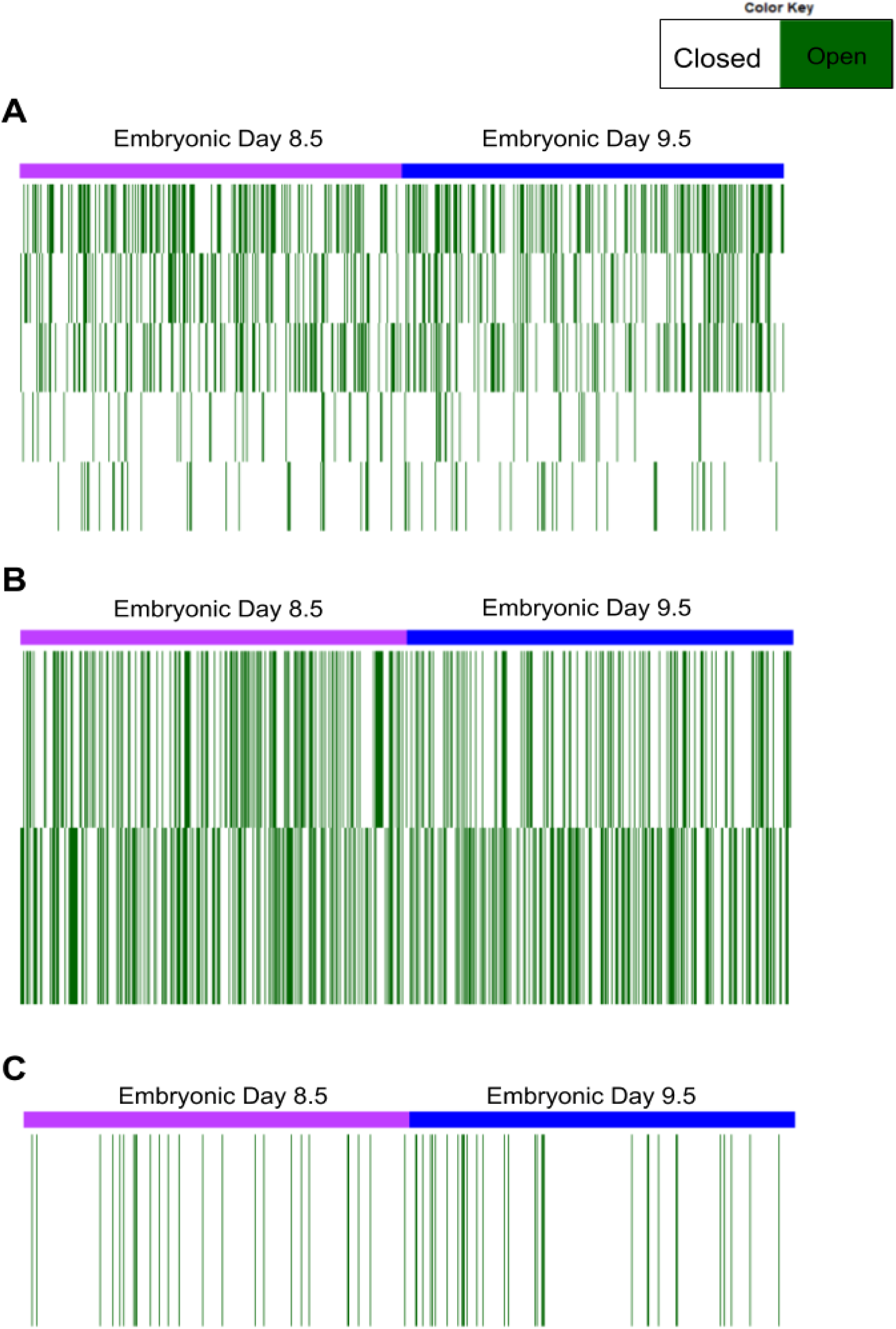
Plot of peaks with overlapping accessibility for the **A** Mef2c, **B** Nkx2-5, and **C** Nppa genes in the Jia et al. cardiac progenitor data.

## Supplemental Tables

**Supplemental Table 1.**
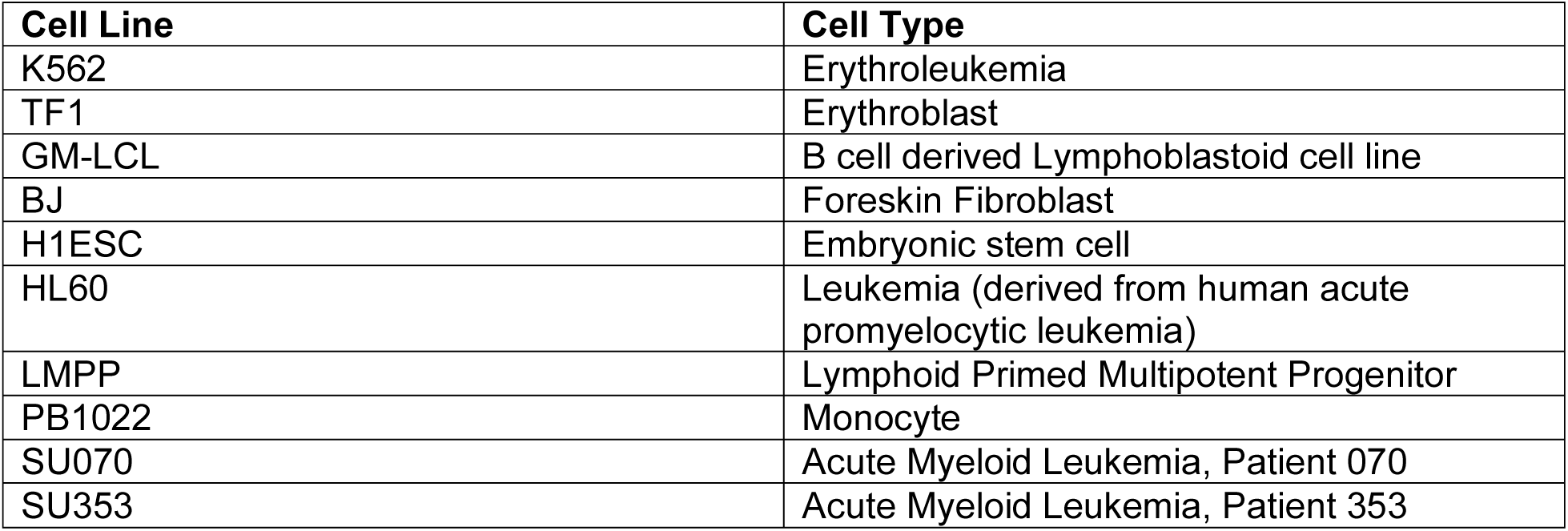
List of the cell lines used in the Schep et al. data set, including the corresponding acronyms used to describe and label them.

| Cell Line | Cell Type |
| --- | --- |
| K562 | Erythroleukemia |
| TF1 | Erythroblast |
| GM-LCL | B cell derived Lymphoblastoid cell line |
| BJ | Foreskin Fibroblast |
| H1ESC | Embryonic stem cell |
| HL60 | Leukemia (derived from human acute promyelocytic leukemia) |
| LMPP | Lymphoid Primed Multipotent Progenitor |
| PB1022 | Monocyte |
| SU070 | Acute Myeloid Leukemia, Patient 070 |
| SU353 | Acute Myeloid Leukemia, Patient 353 |

**Supplemental Table 2.** Annotations of patterns to cell types and the area under the receiver operating curve for these correspondences based on PatternMarker pattern assignment of each cell.

| Pattern | Corresponding Cell Line(s) | AUC |
| --- | --- | --- |
| 1 | K562 Erythroleukemia | 1.00 |
| 2 | TF1 Erythroblast | 1.00 |
| 3 | GM-LCL | 0.996 |
| 4 | BJ Fibroblast | 0.999 |
| 5 | H1ESC | 0.999 |
| 6 | HL60 Leukemia | 0.505 |
| 7 | PB1022, LMPP, SU070, SU353 | 0.86 |

**Supplemental Table 3.**
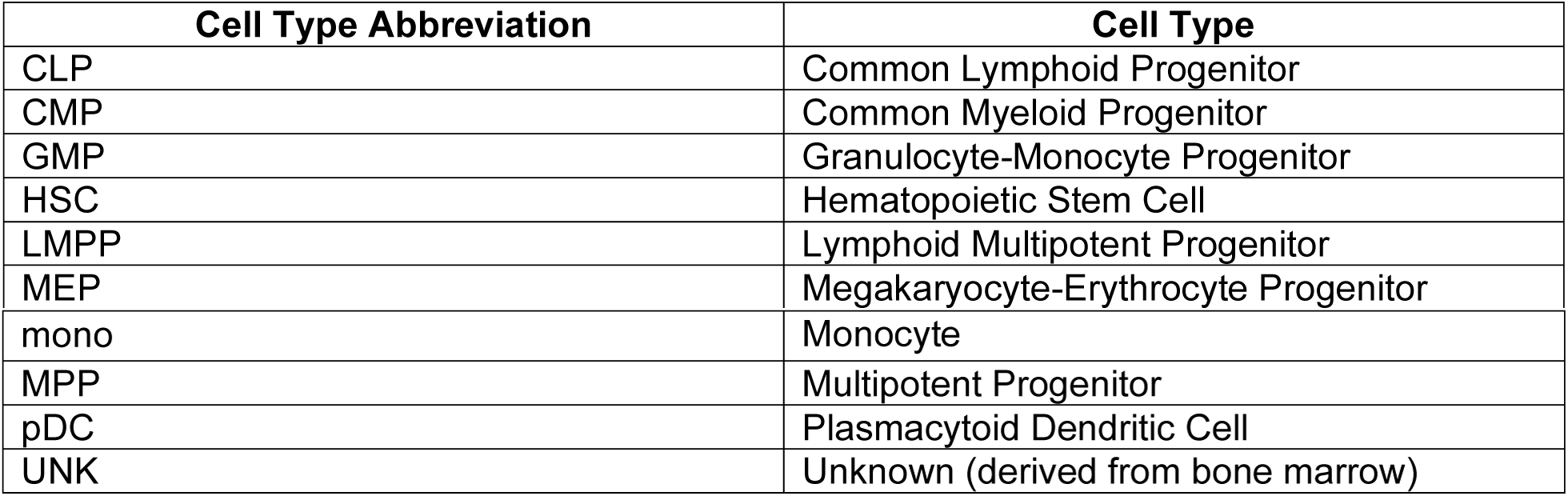
List of the cell lines used in the Buenrostro et al. data set, including the corresponding acronyms used to describe and label them.

| Cell Type Abbreviation | Cell Type |
| --- | --- |
| CLP | Common Lymphoid Progenitor |
| CMP | Common Myeloid Progenitor |
| GMP | Granulocyte-Monocyte Progenitor |
| HSC | Hematopoietic Stem Cell |
| LMPP | Lymphoid Multipotent Progenitor |
| MEP | Megakaryocyte-Erythrocyte Progenitor |
| mono | Monocyte |
| MPP | Multipotent Progenitor |
| pDC | Plasmacytoid Dendritic Cell |
| UNK | Unknown (derived from bone marrow) |

