## Supplemental File 1 for "Matrix factorization and transfer learning uncover regulatory biology across multiple single-cell ATAC-seq data sets"

### CoGAPS Identified Pathways

```
## $`Pattern 1`
## $`Pattern 1`$summaryTable
##                             pathway       PValue
## 5         HALLMARK_HEME_METABOLISM 1.348010e-16
## 6         HALLMARK_MITOTIC_SPINDLE 8.585937e-07
## 8 HALLMARK_TNFA_SIGNALING_VIA_NFKB 3.155078e-05
## 3               HALLMARK_APOPTOSIS 4.977247e-05
## 1            HALLMARK_ADIPOGENESIS 5.795468e-05
## 4 HALLMARK_ESTROGEN_RESPONSE_EARLY 5.795468e-05
## 7      HALLMARK_TGF_BETA_SIGNALING 7.693160e-05
## 2         HALLMARK_APICAL_JUNCTION 1.043982e-04
##
##
## $`Pattern 2`
## $`Pattern 2`$summaryTable
##                              pathway       PValue
## 12          HALLMARK_UV_RESPONSE_DN 5.798672e-12
## 6      HALLMARK_IL2_STAT5_SIGNALING 9.539725e-12
## 4          HALLMARK_HEME_METABOLISM 6.511603e-11
## 9         HALLMARK_MTORC1_SIGNALING 1.284655e-08
## 5                  HALLMARK_HYPOXIA 2.936508e-07
## 8          HALLMARK_MITOTIC_SPINDLE 9.333264e-07
## 1             HALLMARK_ADIPOGENESIS 2.501268e-06
## 2        HALLMARK_ANDROGEN_RESPONSE 1.427122e-05
## 11 HALLMARK_TNFA_SIGNALING_VIA_NFKB 1.789366e-05
## 10       HALLMARK_PROTEIN_SECRETION 1.821590e-05
## 7    HALLMARK_INFLAMMATORY_RESPONSE 6.032164e-05
## 3  HALLMARK_ESTROGEN_RESPONSE_EARLY 1.076035e-04
##
##
## $`Pattern 3`
## $`Pattern 3`$summaryTable
##                                pathway       PValue
## 15   HALLMARK_TNFA_SIGNALING_VIA_NFKB 3.868026e-18
## 7        HALLMARK_IL2_STAT5_SIGNALING 5.130824e-16
## 16            HALLMARK_UV_RESPONSE_DN 4.574162e-14
## 9      HALLMARK_INFLAMMATORY_RESPONSE 8.532207e-12
## 11         HALLMARK_KRAS_SIGNALING_UP 2.259654e-11
## 1        HALLMARK_ALLOGRAFT_REJECTION 1.489475e-10
## 10 HALLMARK_INTERFERON_GAMMA_RESPONSE 1.489475e-10
## 3                  HALLMARK_APOPTOSIS 3.766399e-08
## 8    HALLMARK_IL6_JAK_STAT3_SIGNALING 4.507967e-08
## 5    HALLMARK_ESTROGEN_RESPONSE_EARLY 4.381401e-06
## 12           HALLMARK_MITOTIC_SPINDLE 1.195938e-05
## 2            HALLMARK_APICAL_JUNCTION 1.587116e-05
## 14        HALLMARK_TGF_BETA_SIGNALING 5.198330e-05
## 13          HALLMARK_MTORC1_SIGNALING 5.321764e-05
## 4                 HALLMARK_COMPLEMENT 9.468292e-05
## 6                    HALLMARK_HYPOXIA 2.830334e-04
##
##
```

```
## $`Pattern 4`
## $`Pattern 4`$summaryTable
##                                        pathway       PValue
## 6  HALLMARK_EPITHELIAL_MESENCHYMAL_TRANSITION 8.529163e-36
## 16                     HALLMARK_UV_RESPONSE_DN 1.051232e-25
## 15           HALLMARK_TNFA_SIGNALING_VIA_NFKB 3.232202e-15
## 9                HALLMARK_IL2_STAT5_SIGNALING 3.003042e-14
## 8                            HALLMARK_HYPOXIA 2.573234e-13
## 7            HALLMARK_ESTROGEN_RESPONSE_EARLY 7.308530e-13
## 3                    HALLMARK_APICAL_JUNCTION 2.523192e-10
## 13                        HALLMARK_MYOGENESIS 3.588584e-09
## 4                          HALLMARK_APOPTOSIS 8.742780e-09
## 12                   HALLMARK_MITOTIC_SPINDLE 1.372606e-08
## 5                         HALLMARK_COMPLEMENT 8.851363e-07
## 10            HALLMARK_INFLAMMATORY_RESPONSE 8.851363e-07
## 2                       HALLMARK_ANGIOGENESIS 2.587624e-06
## 11                 HALLMARK_KRAS_SIGNALING_UP 6.999488e-06
## 14                 HALLMARK_TGF_BETA_SIGNALING 5.751367e-05
## 1                  HALLMARK_ANDROGEN_RESPONSE 1.415445e-04
##
##
## $`Pattern 5`
## $`Pattern 5`$summaryTable
##                                        pathway       PValue
## 1                    HALLMARK_APICAL_JUNCTION 1.388378e-13
## 2  HALLMARK_EPITHELIAL_MESENCHYMAL_TRANSITION 3.301153e-12
## 3            HALLMARK_ESTROGEN_RESPONSE_EARLY 2.436182e-11
## 10                     HALLMARK_UV_RESPONSE_DN 3.808226e-10
## 9            HALLMARK_TNFA_SIGNALING_VIA_NFKB 5.766965e-09
## 6                HALLMARK_IL2_STAT5_SIGNALING 3.051722e-07
## 8                         HALLMARK_MYOGENESIS 1.291245e-06
## 7                  HALLMARK_KRAS_SIGNALING_UP 2.576170e-06
## 5                            HALLMARK_HYPOXIA 5.036962e-06
## 4                  HALLMARK_HEDGEHOG_SIGNALING 1.402250e-04
##
##
## $`Pattern 6`
## $`Pattern 6`$summaryTable
##                                        pathway       PValue
## 20           HALLMARK_TNFA_SIGNALING_VIA_NFKB 3.085790e-13
## 11            HALLMARK_INFLAMMATORY_RESPONSE 5.627103e-10
## 10               HALLMARK_IL2_STAT5_SIGNALING 1.693213e-09
## 13                   HALLMARK_MITOTIC_SPINDLE 2.173996e-09
## 12                 HALLMARK_KRAS_SIGNALING_UP 3.659294e-08
## 14                  HALLMARK_MTORC1_SIGNALING 9.468201e-08
## 21                     HALLMARK_UV_RESPONSE_DN 1.216962e-07
## 5            HALLMARK_ESTROGEN_RESPONSE_EARLY 5.707008e-07
## 6                      HALLMARK_G2M_CHECKPOINT 5.707008e-07
## 19                 HALLMARK_TGF_BETA_SIGNALING 1.275552e-06
## 18           HALLMARK_PI3K_AKT_MTOR_SIGNALING 3.213048e-06
## 1                       HALLMARK_ADIPOGENESIS 6.576505e-06
## 4  HALLMARK_EPITHELIAL_MESENCHYMAL_TRANSITION 6.576505e-06
## 17                        HALLMARK_P53_PATHWAY 6.576505e-06
## 3                         HALLMARK_E2F_TARGETS 5.713666e-05
```

```
## 22        HALLMARK_WNT_BETA_CATENIN_SIGNALING 7.632301e-05
## 15                    HALLMARK_MYC_TARGETS_V1 1.106565e-04
## 2                          HALLMARK_APOPTOSIS 1.450926e-04
## 7                         HALLMARK_GLYCOLYSIS 2.082222e-04
## 9                            HALLMARK_HYPOXIA 2.082222e-04
## 8                    HALLMARK_HEME_METABOLISM 3.809065e-04
## 16                        HALLMARK_MYOGENESIS 3.809065e-04
##
##
## $`Pattern 7`
## $`Pattern 7`$summaryTable
##                              pathway       PValue
## 17   HALLMARK_TNFA_SIGNALING_VIA_NFKB 9.869770e-27
## 10     HALLMARK_INFLAMMATORY_RESPONSE 4.117782e-22
## 8        HALLMARK_IL2_STAT5_SIGNALING 1.122470e-09
## 18            HALLMARK_UV_RESPONSE_DN 1.417200e-09
## 11 HALLMARK_INTERFERON_GAMMA_RESPONSE 1.422270e-08
## 7                    HALLMARK_HYPOXIA 3.184648e-08
## 3                  HALLMARK_APOPTOSIS 9.020828e-08
## 16        HALLMARK_TGF_BETA_SIGNALING 1.071393e-07
## 12         HALLMARK_KRAS_SIGNALING_UP 1.503725e-07
## 13           HALLMARK_MITOTIC_SPINDLE 4.678593e-07
## 4                 HALLMARK_COMPLEMENT 6.558180e-07
## 5    HALLMARK_ESTROGEN_RESPONSE_EARLY 6.558180e-07
## 1        HALLMARK_ALLOGRAFT_REJECTION 1.329669e-06
## 9    HALLMARK_IL6_JAK_STAT3_SIGNALING 2.381951e-06
## 15               HALLMARK_P53_PATHWAY 5.153906e-06
## 2          HALLMARK_ANDROGEN_RESPONSE 9.740774e-06
## 19            HALLMARK_UV_RESPONSE_UP 5.921498e-05
## 6     HALLMARK_ESTROGEN_RESPONSE_LATE 3.212447e-04
## 14          HALLMARK_MTORC1_SIGNALING 3.212447e-04
```
