## Supplemental File 2 for "Matrix factorization and transfer learning uncover regulatory biology across multiple single-cell ATAC-seq data sets"

### TF Enrichment

```
## $`Pattern 1`
##              (Other)               ZNF263           GATA1::TAL1
##                  961                  801                   404
##                 EGR1                  SP2                  CTCF
##                  379                  378                   296
##                RREB1             MAF::NFE2                   SP1
##                  211                  184                   175
##                 KLF5                  SP4                 FOXP1
##                  153                  144                   135
##                 IRF1                GATA2                 FOSL1
##                  132                  110                   109
##                STAT1                 NFE2                  E2F6
##                   98                   94                    86
##                 JUND                PLAG1                 GATA3
##                   78                   77                    74
##                NR2C2 SMAD2::SMAD3::SMAD4                 FOSL2
##                   72                   71                    68
##           NFIC::TLX1                KLF16                 HNF4G
##                   65                   62                    61
##                 PAX5           JUN(var.2)                  NFYA
##                   61                   60                    59
##               ZBTB18                  YY1                 ESRRB
##                   59                   57                    55
##                 ZEB1                TBX15                 STAT3
##                   55                   54                    53
##                PRDM1                INSM1                  REST
##                   52                   51                    51
##                 ESR2               POU2F2          STAT1::STAT2
##                   48                   48                    48
##           RARA::RXRA                  SP8                 MEF2C
##                   46                   45                    44
##               SREBF1                NR2F1                  NRF1
##                   44                   43                    42
##        TFAP2B(var.2)                 USF2                  SPIC
##                   41                   41                    37
##            BATF::JUN                 CDX2           JUND(var.2)
##                   36                   35                    35
##                 LEF1                 NFYB                 NHLH1
##                   35                   35                    35
##                 RELA               SREBF2             TAL1::TCF3
##                   35                   34                    34
##                  JUN                KLF14                  ZIC3
##                   33                   33                    33
##                  MSC               ZNF740                  JDP2
##                   32                   32                    31
##                 MAFK                FOXA1                   FOS
##                   31                   30                    28
##                 EBF1           EWSR1-FLI1                POU1F1
##                   27                   26                    25
##                 MAFF        TFAP2B(var.3)                 CEBPA
##                   23                   23                    22
```

```
##               MEF2A               RUNX3       TFAP2A(var.3)
##                  22                  22                  22
##                USF1               TFAP4                DUX4
##                  22                  21                  20
##                ELK4            MAX::MYC         RORA(var.2)
##                  20                  20                  20
##              TCF7L2               TGIF2                E2F4
##                  20                  20                  19
##               FOXB1               FOXH1       TFAP2C(var.3)
##                  19                  19                  19
##                ZIC4               NFKB2                JUNB
##                  19                  18                  17
##         RARA(var.2)                SPI1                ZIC1
##                  17                  17                  17
##               FOXP2                ATF4              HOXB13
##                  16                  15                  15
##                HSF2
##                  15
##
## $`Pattern 2`
##             (Other)                IRF1              ZNF263
##                1219                 592                 497
##               FOXP1          GATA1::TAL1               GATA2
##                 487                 460                 265
##               MEF2C                SPIC                CTCF
##                 263                 231                 228
##                SPI1               GATA3         STAT1::STAT2
##                 189                 182                 166
##               STAT1           MAF::NFE2          JUN(var.2)
##                 163                 157                 152
##              POU2F2               PRDM1                NFE2
##                 152                 146                 141
##                CDX2            BATF::JUN               FOXC2
##                 123                 113                 101
##               FOXA1               RREB1                JUND
##                  98                  97                  96
##                DUX4              POU1F1               RUNX3
##                  86                  86                  81
##               FOXB1               MEF2A               FOSL1
##                  80                  78                  77
##              ZBTB18         JUND(var.2)                 JUN
##                  77                  73                  71
##                MAFK              POU4F3                 FOS
##                  71                  68                  66
##              TCF7L2             ONECUT3              POU4F1
##                  66                  64                  64
##              POU3F3               CEBPA               ESRRB
##                  63                  62                  62
##               HNF1A                JDP2                NFYA
##                  61                  61                  59
##               PROP1               STAT3               FOXP2
##                  59                  58                  57
##               FOXF2 SMAD2::SMAD3::SMAD4                LEF1
##                  55                  54                  53
```

```
##            HOXB13          RORA(var.2)                 MAFF
##                49                   49                   45
##              NFYB           EWSR1-FLI1                HNF4G
##                45                   44                   44
##             NFIL3                 EGR1                  YY1
##                44                   43                   43
##             FOXH1           TAL1::TCF3                FOSL2
##                42                   41                   40
##               MSC               POU3F4           RARA::RXRA
##                40                   40                   40
##        NFIC::TLX1                NR2F1                FOXO3
##                38                   38                   37
##              PAX5                 USF2                 IRF2
##                37                   37                   35
##              PBX1                 HSF2                NR2C2
##                35                   33                   33
##            POU3F2               POU4F2               POU6F2
##                33                   33                   33
##              MAFG              BHLHE23                 DUXA
##                32                   31                   31
##              IRF7               PHOX2A               HOXD13
##                31                   30                   29
##              HSF4                  TEF                 RELA
##                27                   26                   25
##             TGIF2                FOXC1                 REST
##                25                   24                   24
##             TBX15                 ETV2                MEF2B
##                24                   23                   23
##               SP2                  SP1                 MYF6
##                23                   22                   21
##              ATF4                 ESR2                RUNX2
##                20                   20                   20
##               SRF
##                20
##
## $`Pattern 3`
##           (Other)                 IRF1               ZNF263
##              1940                 1215                  735
##       STAT1::STAT2                FOXP1                MEF2C
##               541                  501                  369
##            POU2F2                PRDM1                 CTCF
##               311                  282                  252
##             RREB1            BATF::JUN                 RELA
##               216                  208                  193
##             HNF1A           JUN(var.2)                 EGR1
##               189                  182                  178
##              NFE2            MAF::NFE2                 SPIC
##               161                  154                  150
##              SPI1                 CDX2                  SP2
##               141                  118                  118
##             MEF2A                 PAX5                TBX15
##               116                  115                  107
##              JDP2               TCF7L2               POU1F1
##                95                   94                   91
```

```
##               HNF4G                LEF1               STAT1
##                  89                  89                  89
##              ZBTB18                IRF2              POU4F1
##                  89                  86                  86
##               FOXB1                DUX4                EBF1
##                  85                  82                  82
##                IRF7               PROP1 SMAD2::SMAD3::SMAD4
##                  81                  81                  79
##               GATA2               FOXC2               ESRRB
##                  77                  76                  75
##         RORA(var.2)         JUND(var.2)                 JUN
##                  74                  73                  72
##                JUND               NFKB2              POU3F3
##                  72                  72                  72
##               RUNX3               HNF1B                 FOS
##                  72                  68                  67
##               FOXA1         GATA1::TAL1               FOSL1
##                  67                  67                  64
##                MAFK               STAT3                NFYA
##                  64                  63                  61
##              POU4F3               CEBPA               FOXP2
##                  60                  59                  58
##                ZEB1                 MSC                 YY1
##                  58                  56                  56
##               NFIL3          TAL1::TCF3               GATA3
##                  55                  55                  54
##              POU3F4                 SP1             ONECUT3
##                  53                  53                  48
##                NFYB                TBR1               FOXH1
##                  47                  47                  45
##                MAFF               MEF2D              POU3F2
##                  45                  45                  44
##                USF2               NR2C2                ESR2
##                  44                  43                  42
##                KLF5               MEF2B               FOXO3
##                  42                  42                  41
##               INSM1                IRF8               NR2F1
##                  41                  41                  41
##                PBX1          RARA::RXRA              POU4F2
##                  41                  41                  40
##                 SP4               TBX20               TBX21
##                  40                  40                  40
##          EWSR1-FLI1              HOXB13                NRF1
##                  39                  39                  39
##                ELF3               FOSL2          NFIC::TLX1
##                  38                  38                  37
##                MYF6               EOMES              POU6F2
##                  36                  35                  35
##                E2F6
##                  34
##
## $`Pattern 4`
##             (Other)              ZNF263               FOXP1
##                1210                 541                 386
```

```
##                IRF1            JUN(var.2)                  NFE2
##                 327                   325                   264
##            MAF::NFE2                  JUND                 MEF2C
##                 244                   241                   232
##             BATF::JUN                 FOSL1                   FOS
##                 223                   222                   211
##                FOSL2                POU2F2                 STAT1
##                 156                   122                   122
##                RREB1                  JDP2          STAT1::STAT2
##                 118                   116                   114
##                 CDX2                 PRDM1                   JUN
##                 113                   107                   100
##                 EGR1                 MEF2A                 GATA2
##                  98                    94                    90
##                FOXA1                 FOXB1                 FOXC2
##                  88                    88                    88
##               ZBTB18                 CEBPA           JUND(var.2)
##                  88                    86                    86
##                STAT3                  DUX4                  SPIC
##                  80                    67                    62
##                FOXP2            GATA1::TAL1                 GATA3
##                  59                    59                    57
##               POU4F3                  EBF1                  MAFK
##                  57                    56                    55
##                PROP1                   SP2                 FOXF2
##                  53                    53                    52
##                HNF1A                TCF7L2                 RUNX3
##                  52                    51                    50
##            TAL1::TCF3                  HSF1                 NFIL3
##                  50                    49                    48
##          RORA(var.2)                 FOXH1                NR2C2
##                  48                    47                    47
##               POU4F1 SMAD2::SMAD3::SMAD4        TFAP2B(var.2)
##                  47                    47                    47
##                 RELA                 TBX15                  CTCF
##                  46                    46                    45
##                 LEF1                 ESRRB                 FOXO3
##                  45                    44                    43
##                HNF4G                HOXB13                  ATF4
##                  43                    43                    42
##               POU1F1                  HSF2                POU3F3
##                  40                    39                    39
##            RARA::RXRA                  KLF5                POU4F2
##                  39                    38                    38
##            EWSR1-FLI1                  MAFF                 MEF2B
##                  37                    37                    37
##                 NFYB                  USF2                   MSC
##                  37                    36                    35
##                MEF2D                  E2F6                   SP1
##                  34                    32                    31
##                 PAX5                POU3F2         TFAP2B(var.3)
##                  29                    29                    29
##            NFIC::TLX1                  ESR2                  JUNB
##                  28                    27                    27
```

```
##            PLAG1            POU6F2              RFX3
##               27                27                27
##            TFAP4            HOXC13             INSM1
##               27                26                26
##            NR2F1           ONECUT3              MYF6
##               26                26                25
##            TEAD3               YY1            HOXD13
##               25                25                24
##             HSF4              IRF2            POU3F4
##               24                24                24
##              SRF
##               24
##
## $`Pattern 5`
##             CTCF            ZNF263           (Other)
##             1748              1411               930
##              SP2              EGR1               SP1
##              922               755               294
##            RREB1               SP4              KLF5
##              286               233               191
##             REST              E2F6              NRF1
##              165               143               135
##            PLAG1              ZEB1             NHLH1
##              122                92                85
##            KLF16             INSM1              NFYA
##               83                78                76
##           POU2F2 SMAD2::SMAD3::SMAD4              PAX5
##               73                72                67
##            NR2C2     TFAP2B(var.2)              IRF1
##               66                64                61
##            STAT1             FOXP1              NFYB
##               61                59                58
##    TFAP2B(var.3)              RFX3             KLF14
##               58                56                54
##       EWSR1-FLI1              RELA              ESR2
##               52                51                50
##            PRDM1              RFX5              EBF1
##               50                50                49
##            STAT3             HNF4G              ZIC3
##               49                48                47
##             E2F4       STAT1::STAT2     TFAP2A(var.3)
##               45                43                43
##             ZIC4        NFIC::TLX1             TBX15
##               43                42                40
##           ZBTB18          MAF::NFE2               MSC
##               40                39                39
##            NFKB2     TFAP2A(var.2)              USF2
##               38                37                37
##              YY1            POU1F1     TFAP2C(var.3)
##               37                36                36
##           ZNF740              RFX2               SP8
##               36                34                34
##           SREBF1             ESRRB             FOSL1
##               32                31                30
```

```
##          RARA::RXRA                SPIC               NR2F1
##                  29                  29                  28
##          JUN(var.2)          TAL1::TCF3              TCF7L2
##                  27                  26                  26
##               GLIS2               HINFP                EGR3
##                  25                  25                  24
##               GATA2                 JUN         RORA(var.2)
##                  24                  24                  24
##               FOSL2                LEF1               FOXH1
##                  23                  23                  22
##         GATA1::TAL1              TFAP2B                ZIC1
##                  22                  22                  22
##           BATF::JUN              SREBF2                MAFK
##                  21                  21                  20
##              POU3F4                USF1              ZNF143
##                  20                  20                  20
##                CDX2               FOXP2                GLI2
##                  19                  19                  19
##               TFAP4                JUND                RFX4
##                  19                  18                  18
##                ELK4                 ID4         JUND(var.2)
##                  17                  17                  17
##               KLF13                EGR4                 FOS
##                  17                  16                  16
##            MAX::MYC              ZBTB7A             BHLHE41
##                  16                  16                  15
##                EGR2
##                  15
##
## $`Pattern 6`
##             (Other)              ZNF263                CTCF
##                4404                3142                3019
##                 SP2                EGR1                IRF1
##                1107                1096                 771
##                SPIC               RREB1               FOXP1
##                 648                 616                 480
##                 SP1                SPI1           MAF::NFE2
##                 472                 442                 363
##        STAT1::STAT2                KLF5                NRF1
##                 359                 343                 325
##               PRDM1                E2F6                 SP4
##                 305                 273                 259
##               MEF2C               STAT1              ZBTB18
##                 258                 245                 241
##                PAX5          JUN(var.2)               PLAG1
##                 238                 226                 225
##                NFE2               CEBPA                 YY1
##                 224                 222                 221
## SMAD2::SMAD3::SMAD4         GATA1::TAL1                ZEB1
##                 213                 211                 209
##              POU2F2               ESRRB               TBX15
##                 205                 192                 172
##               NR2C2               FOSL1               INSM1
##                 171                 169                 169
```

```
##              JUND              REST    TFAP2B(var.2)
##               165               163               163
##             STAT3              EBF1             NHLH1
##               156               154               154
##          BATF::JUN        NFIC::TLX1               JUN
##               151               151               143
##              ESR2             KLF16             NR2F1
##               142               141               141
##             HNF4G              RELA             RUNX3
##               140               137               134
##              USF2             GATA2              NFYA
##               133               132               131
##             FOSL2              ATF4             GATA3
##               130               129               125
##               MSC      JUND(var.2)     TFAP2B(var.3)
##               125               121               121
##              CDX2             FOXH1            TCF7L2
##               119               115               115
##              NFYB        EWSR1-FLI1             MEF2A
##               113               111               111
##        RARA::RXRA            ZNF740               FOS
##               109               105               103
##              MAFK              DUX4             FOXA1
##               101               100                97
##             FOXP2        TAL1::TCF3             FOXB1
##                97                96                94
##             KLF14              ELK4            SREBF1
##                93                92                89
##       RORA(var.2)            POU1F1              LEF1
##                88                86                82
##     TFAP2A(var.3)             FOXC2              ZIC3
##                80                79                79
##            SREBF2              MAFF            NFKB2
##                76                75                75
##               SP8              ELK1              RFX5
##                72                71                71
##           MAX::MYC              MYF6              JDP2
##                70                70                69
##            POU4F1           ONECUT3              IRF2
##                69                68                67
##              ZIC1              E2F4            HOXB13
##                67                66                66
##              ELF3
##                65
##
## $`Pattern 7`
##           (Other)            ZNF263              IRF1
##              1620              1063               621
##              CTCF              SPIC               SP2
##               503               371               353
##              EGR1      STAT1::STAT2             FOXP1
##               328               295               240
##              SPI1             RREB1          MAF::NFE2
##               231               208               161
```

```
##                PRDM1                MEF2C                CEBPA 
##                  160                  150                  139 
##                  SP1                  SP4                 NRF1 
##                  128                  125                  115 
##               POU2F2           JUN(var.2)                 KLF5 
##                  113                   96                   96 
##                  JUN                  YY1                FOSL1 
##                   90                   88                   84 
##                STAT1                 ATF4                NR2C2 
##                   80                   76                   76 
##                ESRRB                 CDX2               ZBTB18 
##                   75                   74                   74 
##                HNF4G                TBX15                 NFE2 
##                   72                   72                   70 
##                MEF2A                 NFYA             BATF::JUN 
##                   68                   65                   63 
##                 RELA                 PAX5 SMAD2::SMAD3::SMAD4 
##                   63                   62                   62 
##                FOSL2          JUND(var.2)                 NFYB 
##                   61                   61                   60 
##                 E2F6                 JUND                 USF2 
##                   58                   56                   56 
##                PLAG1                INSM1           EWSR1-FLI1 
##                   55                   53                   52 
##                FOXB1                NR2F1                STAT3 
##                   52                   52                   52 
##                 ZEB1                FOXA1                RUNX3 
##                   50                   49                   48 
##                 MAFF                FOXH1                 MAFK 
##                   47                   46                   46 
##                NFIL3            RARA::RXRA           NFIC::TLX1 
##                   46                   46                   45 
##                 REST               TCF7L2                 DUX4 
##                   45                   45                   44 
##                  FOS                FOXP2                GATA2 
##                   43                   41                   41 
##          RORA(var.2)                  TEF                 ESR2 
##                   41                   41                   40 
##                FOXC2        TFAP2B(var.2)        TFAP2B(var.3) 
##                   40                   40                   40 
##                NFKB2                NHLH1                 RFX3 
##                   39                   39                   38 
##                  MSC               POU1F1                 LEF1 
##                   37                   36                   35 
##                 HSF4                 IRF2                PROP1 
##                   33                   33                   32 
##                 RFX5            TAL1::TCF3                FOXF2 
##                   32                   32                   31 
##          GATA1::TAL1                 HSF1                KLF16 
##                   31                   31                   31 
##                MEF2B               SREBF2                 PBX1 
##                   31                   31                   30 
##               POU4F1                 EBF1                 IRF7 
##                   30                   29                   29 
```

```
##                KLF14              POU4F3               TFAP4 
##                   29                  29                  29 
##                 ELF3              POU3F3               GATA3 
##                   28                  28                  27 
##                 JDP2 
##                   27 
```
