## Supplemental File 3 for "Matrix factorization and transfer learning uncover regulatory biology across multiple single-cell ATAC-seq data sets"

### PatternMarker Motif Overlaps

PatternMarker Motifs Overlapping between Fibroblast Associated Patterns

```
##  [1] "M02896_2.00" "M02913_2.00" "M03027_2.00" "M03170_2.00" "M03354_2.00"
##  [6] "M03370_2.00" "M03505_2.00" "M03507_2.00" "M03540_2.00" "M03554_2.00"
## [11] "M03573_2.00" "M03807_2.00" "M04248_2.00" "M04507_2.00" "M04570_2.00"
## [16] "M04649_2.00" "M04808_2.00" "M07552_2.00" "M07572_2.00" "M07629_2.00"
## [21] "M07768_2.00" "M08887_2.00" "M08957_2.00" "M10652_2.00" "M10935_2.00"
## [26] "M11138_2.00" "M11197_2.00" "M11384_2.00"
```

PatternMarker Motifs not Overlapping between Fibroblast Associated Patterns

```
##    [1] "M00060_2.00" "M00111_2.00" "M00141_2.00" "M00148_2.00"
##    [5] "M00164_2.00" "M00172_2.00" "M00192_2.00" "M00195_2.00"
##    [9] "M00201_2.00" "M00211_2.00" "M00214_2.00" "M00225_2.00"
##   [13] "M00234_2.00" "M00235_2.00" "M00239_2.00" "M00240_2.00"
##   [17] "M00248_2.00" "M00250_2.00" "M00251_2.00" "M00252_2.00"
##   [21] "M00253_2.00" "M00257_2.00" "M00258_2.00" "M00260_2.00"
##   [25] "M00279_2.00" "M00285_2.00" "M00293_2.00" "M00312_2.00"
##   [29] "M00317_2.00" "M00327_2.00" "M00342_2.00" "M00356_2.00"
##   [33] "M00368_2.00" "M00371_2.00" "M00377_2.00" "M00382_2.00"
##   [37] "M00387_2.00" "M00389_2.00" "M00390_2.00" "M00391_2.00"
##   [41] "M00410_2.00" "M00411_2.00" "M00423_2.00" "M00453_2.00"
##   [45] "M00461_2.00" "M00474_2.00" "M00476_2.00" "M00479_2.00"
##   [49] "M00480_2.00" "M00491_2.00" "M00504_2.00" "M00505_2.00"
##   [53] "M00508_2.00" "M00517_2.00" "M00524_2.00" "M00530_2.00"
##   [57] "M00532_2.00" "M00538_2.00" "M00539_2.00" "M00554_2.00"
##   [61] "M00665_2.00" "M00710_2.00" "M00712_2.00" "M00740_2.00"
##   [65] "M00759_2.00" "M00791_2.00" "M00796_2.00" "M00798_2.00"
##   [69] "M00806_2.00" "M00812_2.00" "M00813_2.00" "M00821_2.00"
##   [73] "M00839_2.00" "M00987_2.00" "M01000_2.00" "M01104_2.00"
##   [77] "M01228_2.00" "M01242_2.00" "M01245_2.00" "M01259_2.00"
##   [81] "M01272_2.00" "M01297_2.00" "M01512_2.00" "M01659_2.00"
##   [85] "M01690_2.00" "M01718_2.00" "M01720_2.00" "M01729_2.00"
##   [89] "M01732_2.00" "M01818_2.00" "M01903_2.00" "M01931_2.00"
##   [93] "M01977_2.00" "M01982_2.00" "M01983_2.00" "M01984_2.00"
##   [97] "M01988_2.00" "M01989_2.00" "M02022_2.00" "M02062_2.00"
##  [101] "M02092_2.00" "M02124_2.00" "M02165_2.00" "M02190_2.00"
##  [105] "M02210_2.00" "M02214_2.00" "M02221_2.00" "M02228_2.00"
##  [109] "M02230_2.00" "M02247_2.00" "M02248_2.00" "M02249_2.00"
##  [113] "M02253_2.00" "M02282_2.00" "M02405_2.00" "M02408_2.00"
##  [117] "M02430_2.00" "M02440_2.00" "M02441_2.00" "M02442_2.00"
##  [121] "M02443_2.00" "M02444_2.00" "M02446_2.00" "M02488_2.00"
##  [125] "M02532_2.00" "M02652_2.00" "M02667_2.00" "M02668_2.00"
##  [129] "M02689_2.00" "M02693_2.00" "M02695_2.00" "M02696_2.00"
##  [133] "M02717_2.00" "M02729_2.00" "M02736_2.00" "M02742_2.00"
##  [137] "M02747_2.00" "M02748_2.00" "M02754_2.00" "M02755_2.00"
##  [141] "M02757_2.00" "M02758_2.00" "M02760_2.00" "M02761_2.00"
##  [145] "M02765_2.00" "M02766_2.00" "M02767_2.00" "M02770_2.00"
##  [149] "M02776_2.00" "M02777_2.00" "M02784_2.00" "M02795_2.00"
##  [153] "M02798_2.00" "M02804_2.00" "M02806_2.00" "M02809_2.00"
```

```
##  [157] "M02812_2.00" "M02819_2.00" "M02820_2.00" "M02823_2.00"
##  [161] "M02824_2.00" "M02826_2.00" "M02827_2.00" "M02828_2.00"
##  [165] "M02832_2.00" "M02833_2.00" "M02835_2.00" "M02840_2.00"
##  [169] "M02841_2.00" "M02842_2.00" "M02843_2.00" "M02844_2.00"
##  [173] "M02845_2.00" "M02846_2.00" "M02847_2.00" "M02849_2.00"
##  [177] "M02851_2.00" "M02852_2.00" "M02859_2.00" "M02863_2.00"
##  [181] "M02864_2.00" "M02867_2.00" "M02868_2.00" "M02869_2.00"
##  [185] "M02872_2.00" "M02883_2.00" "M02886_2.00" "M02887_2.00"
##  [189] "M02889_2.00" "M02893_2.00" "M02894_2.00" "M02904_2.00"
##  [193] "M02906_2.00" "M02909_2.00" "M02912_2.00" "M02915_2.00"
##  [197] "M02917_2.00" "M02932_2.00" "M02934_2.00" "M02939_2.00"
##  [201] "M02942_2.00" "M02943_2.00" "M02944_2.00" "M02947_2.00"
##  [205] "M02949_2.00" "M02954_2.00" "M02957_2.00" "M02963_2.00"
##  [209] "M02974_2.00" "M02975_2.00" "M02993_2.00" "M02995_2.00"
##  [213] "M03000_2.00" "M03008_2.00" "M03009_2.00" "M03028_2.00"
##  [217] "M03038_2.00" "M03039_2.00" "M03042_2.00" "M03051_2.00"
##  [221] "M03061_2.00" "M03069_2.00" "M03070_2.00" "M03085_2.00"
##  [225] "M03086_2.00" "M03089_2.00" "M03091_2.00" "M03101_2.00"
##  [229] "M03110_2.00" "M03122_2.00" "M03127_2.00" "M03130_2.00"
##  [233] "M03132_2.00" "M03134_2.00" "M03140_2.00" "M03158_2.00"
##  [237] "M03162_2.00" "M03166_2.00" "M03176_2.00" "M03184_2.00"
##  [241] "M03202_2.00" "M03217_2.00" "M03224_2.00" "M03228_2.00"
##  [245] "M03229_2.00" "M03236_2.00" "M03242_2.00" "M03249_2.00"
##  [249] "M03252_2.00" "M03253_2.00" "M03255_2.00" "M03266_2.00"
##  [253] "M03274_2.00" "M03278_2.00" "M03283_2.00" "M03303_2.00"
##  [257] "M03316_2.00" "M03322_2.00" "M03325_2.00" "M03328_2.00"
##  [261] "M03330_2.00" "M03332_2.00" "M03335_2.00" "M03336_2.00"
##  [265] "M03340_2.00" "M03341_2.00" "M03344_2.00" "M03345_2.00"
##  [269] "M03346_2.00" "M03347_2.00" "M03350_2.00" "M03352_2.00"
##  [273] "M03356_2.00" "M03369_2.00" "M03372_2.00" "M03385_2.00"
##  [277] "M03389_2.00" "M03398_2.00" "M03401_2.00" "M03404_2.00"
##  [281] "M03418_2.00" "M03419_2.00" "M03429_2.00" "M03438_2.00"
##  [285] "M03439_2.00" "M03440_2.00" "M03441_2.00" "M03451_2.00"
##  [289] "M03473_2.00" "M03477_2.00" "M03478_2.00" "M03481_2.00"
##  [293] "M03482_2.00" "M03483_2.00" "M03484_2.00" "M03485_2.00"
##  [297] "M03488_2.00" "M03489_2.00" "M03491_2.00" "M03494_2.00"
##  [301] "M03495_2.00" "M03496_2.00" "M03498_2.00" "M03499_2.00"
##  [305] "M03503_2.00" "M03509_2.00" "M03516_2.00" "M03517_2.00"
##  [309] "M03518_2.00" "M03521_2.00" "M03523_2.00" "M03526_2.00"
##  [313] "M03530_2.00" "M03535_2.00" "M03537_2.00" "M03538_2.00"
##  [317] "M03546_2.00" "M03551_2.00" "M03565_2.00" "M03566_2.00"
##  [321] "M03569_2.00" "M03577_2.00" "M03579_2.00" "M03584_2.00"
##  [325] "M03586_2.00" "M03587_2.00" "M03588_2.00" "M03589_2.00"
##  [329] "M03595_2.00" "M03600_2.00" "M03602_2.00" "M03605_2.00"
##  [333] "M03639_2.00" "M03640_2.00" "M03641_2.00" "M03655_2.00"
##  [337] "M03656_2.00" "M03666_2.00" "M03673_2.00" "M03676_2.00"
##  [341] "M03677_2.00" "M03716_2.00" "M03727_2.00" "M03736_2.00"
##  [345] "M03737_2.00" "M03745_2.00" "M03746_2.00" "M03747_2.00"
##  [349] "M03748_2.00" "M03764_2.00" "M03765_2.00" "M03767_2.00"
##  [353] "M03768_2.00" "M03781_2.00" "M03784_2.00" "M03801_2.00"
##  [357] "M03803_2.00" "M03812_2.00" "M03828_2.00" "M03841_2.00"
##  [361] "M03850_2.00" "M03855_2.00" "M03856_2.00" "M03862_2.00"
##  [365] "M03867_2.00" "M03877_2.00" "M03886_2.00" "M03892_2.00"
##  [369] "M03893_2.00" "M03894_2.00" "M03899_2.00" "M03900_2.00"
```

```
## [373] "M03909_2.00" "M03924_2.00" "M03926_2.00" "M03933_2.00"
## [377] "M03938_2.00" "M03944_2.00" "M03945_2.00" "M03947_2.00"
## [381] "M03955_2.00" "M03966_2.00" "M03973_2.00" "M03975_2.00"
## [385] "M03978_2.00" "M03979_2.00" "M03981_2.00" "M03982_2.00"
## [389] "M03983_2.00" "M03987_2.00" "M03989_2.00" "M03996_2.00"
## [393] "M04012_2.00" "M04029_2.00" "M04030_2.00" "M04031_2.00"
## [397] "M04042_2.00" "M04045_2.00" "M04046_2.00" "M04047_2.00"
## [401] "M04049_2.00" "M04052_2.00" "M04053_2.00" "M04054_2.00"
## [405] "M04055_2.00" "M04056_2.00" "M04064_2.00" "M04066_2.00"
## [409] "M04067_2.00" "M04068_2.00" "M04069_2.00" "M04070_2.00"
## [413] "M04071_2.00" "M04072_2.00" "M04073_2.00" "M04077_2.00"
## [417] "M04079_2.00" "M04092_2.00" "M04094_2.00" "M04115_2.00"
## [421] "M04116_2.00" "M04118_2.00" "M04119_2.00" "M04122_2.00"
## [425] "M04130_2.00" "M04140_2.00" "M04141_2.00" "M04142_2.00"
## [429] "M04143_2.00" "M04145_2.00" "M04147_2.00" "M04160_2.00"
## [433] "M04161_2.00" "M04162_2.00" "M04166_2.00" "M04168_2.00"
## [437] "M04171_2.00" "M04173_2.00" "M04174_2.00" "M04175_2.00"
## [441] "M04176_2.00" "M04179_2.00" "M04183_2.00" "M04185_2.00"
## [445] "M04187_2.00" "M04189_2.00" "M04204_2.00" "M04225_2.00"
## [449] "M04228_2.00" "M04235_2.00" "M04238_2.00" "M04242_2.00"
## [453] "M04245_2.00" "M04246_2.00" "M04249_2.00" "M04251_2.00"
## [457] "M04252_2.00" "M04254_2.00" "M04260_2.00" "M04266_2.00"
## [461] "M04267_2.00" "M04275_2.00" "M04277_2.00" "M04278_2.00"
## [465] "M04280_2.00" "M04283_2.00" "M04284_2.00" "M04285_2.00"
## [469] "M04287_2.00" "M04289_2.00" "M04291_2.00" "M04302_2.00"
## [473] "M04304_2.00" "M04306_2.00" "M04309_2.00" "M04311_2.00"
## [477] "M04312_2.00" "M04313_2.00" "M04316_2.00" "M04318_2.00"
## [481] "M04319_2.00" "M04324_2.00" "M04330_2.00" "M04332_2.00"
## [485] "M04333_2.00" "M04336_2.00" "M04341_2.00" "M04342_2.00"
## [489] "M04345_2.00" "M04348_2.00" "M04351_2.00" "M04354_2.00"
## [493] "M04373_2.00" "M04377_2.00" "M04383_2.00" "M04384_2.00"
## [497] "M04385_2.00" "M04396_2.00" "M04402_2.00" "M04403_2.00"
## [501] "M04406_2.00" "M04417_2.00" "M04422_2.00" "M04427_2.00"
## [505] "M04447_2.00" "M04450_2.00" "M04454_2.00" "M04459_2.00"
## [509] "M04460_2.00" "M04471_2.00" "M04472_2.00" "M04473_2.00"
## [513] "M04475_2.00" "M04477_2.00" "M04480_2.00" "M04481_2.00"
## [517] "M04482_2.00" "M04485_2.00" "M04493_2.00" "M04502_2.00"
## [521] "M04503_2.00" "M04521_2.00" "M04524_2.00" "M04533_2.00"
## [525] "M04534_2.00" "M04535_2.00" "M04543_2.00" "M04547_2.00"
## [529] "M04549_2.00" "M04551_2.00" "M04564_2.00" "M04565_2.00"
## [533] "M04569_2.00" "M04571_2.00" "M04577_2.00" "M04586_2.00"
## [537] "M04592_2.00" "M04593_2.00" "M04597_2.00" "M04599_2.00"
## [541] "M04600_2.00" "M04601_2.00" "M04603_2.00" "M04604_2.00"
## [545] "M04610_2.00" "M04612_2.00" "M04617_2.00" "M04625_2.00"
## [549] "M04626_2.00" "M04633_2.00" "M04634_2.00" "M04636_2.00"
## [553] "M04651_2.00" "M04653_2.00" "M04664_2.00" "M04665_2.00"
## [557] "M04667_2.00" "M04673_2.00" "M04675_2.00" "M04684_2.00"
## [561] "M04692_2.00" "M04693_2.00" "M04694_2.00" "M04695_2.00"
## [565] "M04697_2.00" "M04699_2.00" "M04707_2.00" "M04729_2.00"
## [569] "M04731_2.00" "M04734_2.00" "M04736_2.00" "M04738_2.00"
## [573] "M04749_2.00" "M04762_2.00" "M04765_2.00" "M04771_2.00"
## [577] "M04773_2.00" "M04778_2.00" "M04781_2.00" "M04785_2.00"
## [581] "M04797_2.00" "M04810_2.00" "M04816_2.00" "M04817_2.00"
## [585] "M04818_2.00" "M04819_2.00" "M04820_2.00" "M04821_2.00"
```

```
## [589] "M04823_2.00" "M04825_2.00" "M04827_2.00" "M04831_2.00"
## [593] "M04833_2.00" "M04836_2.00" "M04842_2.00" "M04844_2.00"
## [597] "M04851_2.00" "M04852_2.00" "M04853_2.00" "M04864_2.00"
## [601] "M04866_2.00" "M04885_2.00" "M04886_2.00" "M04890_2.00"
## [605] "M04900_2.00" "M04935_2.00" "M05010_2.00" "M05011_2.00"
## [609] "M05012_2.00" "M05013_2.00" "M05083_2.00" "M05085_2.00"
## [613] "M05095_2.00" "M05104_2.00" "M05114_2.00" "M05117_2.00"
## [617] "M05119_2.00" "M05152_2.00" "M05159_2.00" "M05233_2.00"
## [621] "M05249_2.00" "M05250_2.00" "M05255_2.00" "M05256_2.00"
## [625] "M05283_2.00" "M05285_2.00" "M05288_2.00" "M05290_2.00"
## [629] "M05298_2.00" "M05299_2.00" "M05301_2.00" "M05302_2.00"
## [633] "M05327_2.00" "M05328_2.00" "M05337_2.00" "M05339_2.00"
## [637] "M05341_2.00" "M05352_2.00" "M05353_2.00" "M05374_2.00"
## [641] "M05375_2.00" "M05376_2.00" "M05388_2.00" "M05389_2.00"
## [645] "M05393_2.00" "M05395_2.00" "M05397_2.00" "M05402_2.00"
## [649] "M05403_2.00" "M05412_2.00" "M05414_2.00" "M05415_2.00"
## [653] "M05416_2.00" "M05419_2.00" "M05420_2.00" "M05422_2.00"
## [657] "M05423_2.00" "M05425_2.00" "M05428_2.00" "M05431_2.00"
## [661] "M05435_2.00" "M05439_2.00" "M05441_2.00" "M05443_2.00"
## [665] "M05445_2.00" "M05453_2.00" "M05456_2.00" "M05457_2.00"
## [669] "M05478_2.00" "M05488_2.00" "M05495_2.00" "M05497_2.00"
## [673] "M05499_2.00" "M05502_2.00" "M05509_2.00" "M05512_2.00"
## [677] "M05514_2.00" "M05515_2.00" "M05518_2.00" "M05523_2.00"
## [681] "M05524_2.00" "M05526_2.00" "M05537_2.00" "M05539_2.00"
## [685] "M05541_2.00" "M05543_2.00" "M05553_2.00" "M05554_2.00"
## [689] "M05557_2.00" "M05568_2.00" "M05569_2.00" "M05571_2.00"
## [693] "M05572_2.00" "M05580_2.00" "M05590_2.00" "M05591_2.00"
## [697] "M05605_2.00" "M05607_2.00" "M05610_2.00" "M05618_2.00"
## [701] "M05622_2.00" "M05623_2.00" "M05624_2.00" "M05638_2.00"
## [705] "M05640_2.00" "M05641_2.00" "M05642_2.00" "M05644_2.00"
## [709] "M05646_2.00" "M05648_2.00" "M05650_2.00" "M05662_2.00"
## [713] "M05664_2.00" "M05668_2.00" "M05672_2.00" "M05677_2.00"
## [717] "M05679_2.00" "M05680_2.00" "M05684_2.00" "M05685_2.00"
## [721] "M05687_2.00" "M05688_2.00" "M05693_2.00" "M05694_2.00"
## [725] "M05695_2.00" "M05697_2.00" "M05701_2.00" "M05704_2.00"
## [729] "M05707_2.00" "M05710_2.00" "M05712_2.00" "M05714_2.00"
## [733] "M05734_2.00" "M05736_2.00" "M05737_2.00" "M05738_2.00"
## [737] "M05743_2.00" "M05744_2.00" "M05749_2.00" "M05750_2.00"
## [741] "M05756_2.00" "M05762_2.00" "M05767_2.00" "M05768_2.00"
## [745] "M05770_2.00" "M05810_2.00" "M05812_2.00" "M05813_2.00"
## [749] "M05818_2.00" "M05819_2.00" "M05823_2.00" "M05824_2.00"
## [753] "M05825_2.00" "M05826_2.00" "M05829_2.00" "M05830_2.00"
## [757] "M05851_2.00" "M05937_2.00" "M05955_2.00" "M05956_2.00"
## [761] "M05985_2.00" "M05991_2.00" "M05993_2.00" "M06005_2.00"
## [765] "M06018_2.00" "M06053_2.00" "M06056_2.00" "M06089_2.00"
## [769] "M06128_2.00" "M06129_2.00" "M06140_2.00" "M06183_2.00"
## [773] "M06184_2.00" "M06195_2.00" "M06205_2.00" "M06238_2.00"
## [777] "M06277_2.00" "M06280_2.00" "M06348_2.00" "M06417_2.00"
## [781] "M06428_2.00" "M06436_2.00" "M06448_2.00" "M06474_2.00"
## [785] "M06559_2.00" "M07538_2.00" "M07812_2.00" "M07815_2.00"
## [789] "M07817_2.00" "M07827_2.00" "M07828_2.00" "M07829_2.00"
## [793] "M07831_2.00" "M07833_2.00" "M07834_2.00" "M07835_2.00"
## [797] "M07837_2.00" "M07838_2.00" "M07933_2.00" "M07935_2.00"
## [801] "M07936_2.00" "M07942_2.00" "M07962_2.00" "M07974_2.00"
```

```
##  [805] "M07975_2.00" "M07980_2.00" "M07987_2.00" "M08012_2.00"
##  [809] "M08035_2.00" "M08036_2.00" "M08037_2.00" "M08043_2.00"
##  [813] "M08048_2.00" "M08052_2.00" "M08054_2.00" "M08058_2.00"
##  [817] "M08059_2.00" "M08064_2.00" "M08065_2.00" "M08067_2.00"
##  [821] "M08069_2.00" "M08070_2.00" "M08071_2.00" "M08076_2.00"
##  [825] "M08093_2.00" "M08098_2.00" "M08116_2.00" "M08118_2.00"
##  [829] "M08120_2.00" "M08121_2.00" "M08123_2.00" "M08131_2.00"
##  [833] "M08135_2.00" "M08137_2.00" "M08138_2.00" "M08149_2.00"
##  [837] "M08164_2.00" "M08165_2.00" "M08175_2.00" "M08188_2.00"
##  [841] "M08196_2.00" "M08199_2.00" "M08203_2.00" "M08212_2.00"
##  [845] "M08217_2.00" "M08428_2.00" "M08455_2.00" "M08478_2.00"
##  [849] "M08578_2.00" "M08590_2.00" "M08706_2.00" "M08710_2.00"
##  [853] "M08783_2.00" "M08786_2.00" "M08787_2.00" "M08797_2.00"
##  [857] "M08803_2.00" "M08804_2.00" "M08805_2.00" "M08806_2.00"
##  [861] "M08811_2.00" "M08815_2.00" "M08821_2.00" "M08828_2.00"
##  [865] "M08832_2.00" "M08836_2.00" "M08837_2.00" "M08846_2.00"
##  [869] "M08847_2.00" "M08862_2.00" "M08883_2.00" "M08890_2.00"
##  [873] "M08899_2.00" "M08902_2.00" "M08905_2.00" "M08909_2.00"
##  [877] "M08916_2.00" "M08918_2.00" "M08928_2.00" "M08929_2.00"
##  [881] "M08953_2.00" "M08977_2.00" "M08980_2.00" "M08984_2.00"
##  [885] "M09011_2.00" "M09013_2.00" "M09024_2.00" "M09025_2.00"
##  [889] "M09055_2.00" "M09057_2.00" "M09083_2.00" "M09084_2.00"
##  [893] "M09087_2.00" "M09090_2.00" "M09091_2.00" "M09093_2.00"
##  [897] "M09094_2.00" "M09098_2.00" "M09100_2.00" "M09102_2.00"
##  [901] "M09103_2.00" "M09105_2.00" "M09111_2.00" "M09112_2.00"
##  [905] "M09113_2.00" "M09114_2.00" "M09122_2.00" "M09128_2.00"
##  [909] "M09130_2.00" "M09141_2.00" "M09142_2.00" "M09153_2.00"
##  [913] "M09158_2.00" "M09161_2.00" "M09165_2.00" "M09166_2.00"
##  [917] "M09170_2.00" "M09173_2.00" "M09174_2.00" "M09181_2.00"
##  [921] "M09185_2.00" "M09187_2.00" "M09192_2.00" "M09200_2.00"
##  [925] "M09206_2.00" "M09209_2.00" "M09214_2.00" "M09216_2.00"
##  [929] "M09217_2.00" "M09219_2.00" "M09220_2.00" "M09221_2.00"
##  [933] "M09247_2.00" "M09250_2.00" "M09252_2.00" "M09253_2.00"
##  [937] "M09255_2.00" "M09263_2.00" "M09276_2.00" "M09279_2.00"
##  [941] "M09283_2.00" "M09290_2.00" "M09309_2.00" "M09310_2.00"
##  [945] "M09311_2.00" "M09313_2.00" "M09316_2.00" "M09320_2.00"
##  [949] "M09326_2.00" "M09332_2.00" "M09335_2.00" "M09336_2.00"
##  [953] "M09337_2.00" "M09338_2.00" "M09339_2.00" "M09342_2.00"
##  [957] "M09345_2.00" "M09350_2.00" "M09353_2.00" "M09354_2.00"
##  [961] "M09360_2.00" "M09365_2.00" "M09370_2.00" "M09377_2.00"
##  [965] "M09387_2.00" "M09400_2.00" "M09403_2.00" "M09407_2.00"
##  [969] "M09413_2.00" "M09414_2.00" "M09415_2.00" "M09418_2.00"
##  [973] "M09420_2.00" "M09427_2.00" "M09431_2.00" "M09434_2.00"
##  [977] "M09437_2.00" "M09438_2.00" "M09486_2.00" "M09495_2.00"
##  [981] "M09545_2.00" "M09566_2.00" "M09581_2.00" "M09587_2.00"
##  [985] "M09603_2.00" "M09616_2.00" "M09638_2.00" "M09653_2.00"
##  [989] "M09672_2.00" "M09674_2.00" "M09679_2.00" "M09684_2.00"
##  [993] "M09694_2.00" "M09802_2.00" "M09816_2.00" "M09864_2.00"
##  [997] "M09938_2.00" "M09942_2.00" "M09954_2.00" "M09955_2.00"
## [1001] "M10029_2.00" "M10074_2.00" "M10127_2.00" "M10129_2.00"
## [1005] "M10307_2.00" "M10423_2.00" "M10425_2.00" "M10527_2.00"
## [1009] "M10530_2.00" "M10540_2.00" "M10550_2.00" "M10558_2.00"
## [1013] "M10565_2.00" "M10570_2.00" "M10576_2.00" "M10580_2.00"
## [1017] "M10588_2.00" "M10605_2.00" "M10740_2.00" "M10746_2.00"
```

```
## [1021] "M10779_2.00" "M10784_2.00" "M10804_2.00" "M10805_2.00"
## [1025] "M10806_2.00" "M10808_2.00" "M10814_2.00" "M10828_2.00"
## [1029] "M10831_2.00" "M10832_2.00" "M10849_2.00" "M10876_2.00"
## [1033] "M10894_2.00" "M10908_2.00" "M10923_2.00" "M10936_2.00"
## [1037] "M10937_2.00" "M10990_2.00" "M11054_2.00" "M11068_2.00"
## [1041] "M11102_2.00" "M11120_2.00" "M11124_2.00" "M11140_2.00"
## [1045] "M11180_2.00" "M11198_2.00" "M11243_2.00" "M11246_2.00"
## [1049] "M11341_2.00" "M11355_2.00" "M11367_2.00" "M11388_2.00"
```
